# Microglia Stimulate Zebrafish Brain Repair Via a Tumor Necrosis Factor-*α*-Initiated Inflammatory Cascade

**DOI:** 10.1101/2020.10.08.330662

**Authors:** Palsamy Kanagaraj, Jessica Y. Chen, Kaia Skaggs, Yusuf Qadeer, Meghan Connors, Noah Cutler, Joshua Richmond, Vineeth Kommidi, Allison Poles, Danielle Affrunti, Curtis Powell, Daniel Goldman, Jack M. Parent

## Abstract

The adult zebrafish brain, unlike mammals, has a remarkable regenerative capacity. Although inflammation inhibits regeneration in mammals, it is necessary for zebrafish brain repair. Microglia are resident brain immune cells that regulate the inflammatory response. To explore the microglial role in repair, we used liposomal clodronate, colony stimulating factor-1 receptor (*csf1r*) inhibition to ablate microglia and two genetic mutants that lacks microglia during brain injury. We found that microglial ablation inhibited injury-induced neurogenesis and regeneration. Microglial suppression specifically attenuated cell proliferation at the progenitor cell amplification stage of neurogenesis. Notably, the loss of microglia impaired phospho-stat3 (signal transducer and activator of transcription 3) and ß-catenin signaling by dynamic regulation of tumor necrosis factor-a after injury, and the ectopic activation of stat3 and ß-catenin rescued neurogenesis defects caused by microglial loss. Microglial absence leads to neutrophil accumulation, hindering the resolution of inflammation and macrophages are not sufficient for regeneration. These findings reveal specific roles of microglia and inflammatory signaling during zebrafish telencephalic regeneration that should provide strategies to improve mammalian brain repair.

## Introduction

Unlike the limited regenerative capacity of the adult mammalian brain, zebrafish show robust repair after brain injury (Kyritsis et al., 2012; Paredes et al., 2015; Parent, 2003; Skaggs et al., 2014; Sorrells et al., 2018). This restorative capacity is not limited to brain as zebrafish are also used to model tissue regeneration in heart, retina, fin, and spinal cord (Alunni & Bally-Cuif, 2016; Becker & Becker, 2008; Goldman, 2014; Poss et al., 2002; Sehring et al., 2016). Although the reasons underlying the differential brain restorative responses are unclear, one factor likely involves adult neurogenesis. The mammalian central nervous system (CNS) has ongoing adult neurogenesis mainly in two forebrain regions, the subventricular zone (SVZ) and the dentate gyrus (DG) of the hippocampal formation (Götz & Huttner, 2005). The adult neural stem cells (aNSCs) of the SVZ and DG are poorly responsive to distant injury and neurodegenerative diseases, and the increased neurogenesis induced by local injury often does not produce viable neurons that survive in the injury environment (Ming & Song, 2011; Parent et al., 2002; Peretto & Bonfanti, 2014).

In contrast to mammals, zebrafish have many adult neurogenic niches in which aNSCs generate neurons that are recruited by injury, migrate to lesion sites and successfully repair tissue damage (Alunni & Bally-Cuif, 2016; Baumgart et al., 2012; Kizil, Kaslin, et al., 2012; Skaggs et al., 2014). Many studies have shown that ventricular zone radial glia act as aNSCs and produce new neurons constitutively (Kroehne et al., 2011; Skaggs et al., 2014; Than-Trong & Bally-Cuif, 2015). Like the mammalian CNS, zebrafish maintain aNSC quiescence to preserve the NSC pool and the ability to divide upon activation (Alvarez-Buylla & Garcia-Verdugo, 2002; Cheung & Rando, 2013; Gage, 2002; Katz et al., 2016; Pilz et al., 2018). The aNSCs divide asymmetrically to retain or symmetrically to expand the NSC pool during constitutive neurogenesis. Only after injury, however, do aNSCs divide symmetrically to produce two neural progenitor cells (NPCs) that divide again symmetrically to produce 4 daughter cells(Barbosa & Ninkovic, 2016; Barbosa et al., 2015). unique symmetric division after injury amplifies the NPC pool to promote robust regeneration in zebrafish.

Another difference between mammals and zebrafish involves inhibitory factors in the mammalian CNS that block regeneration, such as those derived from reactive glia and non-CNS inflammatory cells (Fitch & Silver, 2008; Gerlach et al., 2016). Zebrafish do not have parenchymal astrocytes that induce glial scarring and express molecules that inhibit brain repair (Baumgart et al., 2012; Ghosh & Hui, 2016; Zambusi & Ninkovic, 2020; Zupanc & Sirbulescu, 2011). The CNS inflammatory response to injury also differs between mammals and zebrafish (Bosak et al., 2018). Mammalian inflammatory responses have been shown to play opposing roles in repair depending upon the injury context, and inflammation inhibits tissue regeneration in multiple CNS injury models (Ekdahl et al., 2003; Fitch & Silver, 2008; Iosif et al., 2006; Monje et al., 2003). In contrast, acute inflammation appears necessary for promoting regeneration after injury in zebrafish. Inflammatory mediators implicated in the zebrafish CNS pro-regenerative response include leukotrienes in brain (Kyritsis et al., 2012), interleukin (il) 6, il11, and leptin in retina (Wan et al., 2014; Zhao et al., 2014), and tnf*α* and Il1*β* in spinal cord (T. M. Tsarouchas et al., 2018).

Inflammation is a complex process involving various immune cell types and multiple signaling molecules secreted or induced by the immune cells. As the resident immune cells of the CNS, microglia rapidly respond to brain injury. When activated, they secrete various cytokines and chemokines to attract other cell types and regulate inflammation (Ekdahl et al., 2009; Jin & Yamashita, 2016). In mammals, microglia take on M1 pro-inflammatory states that are neurotoxic or M2 anti-inflammatory states that are neuroprotective after CNS injury (Crain et al., 2013; Jassam et al., 2017; Ransohoff, 2016). Macrophages are also critical for regeneration of fin, retina, and heart in zebrafish (Huang et al., 2012; Petrie et al., 2014; Sanz-Morejon et al., 2019). In contrast, several studies showed that microglia are not necessary for murine axonal regeneration (Hilla et al., 2017) and spinal cord repair in zebrafish (T. M. Tsarouchas et al., 2018). Though many studies have shown that the microglial response after injury, whether microglia are required for successful repair after acute brain injury in zebrafish is unknown.

Here we show that the microglia are crucial for successful telencephalic injury-induced cell proliferation, neurogenesis, and regeneration in adult zebrafish. We used liposomal clodronate to suppress activated microglia and found decreased proliferation of sox2-expressing intermediate progenitor cells. Two genetic mutants confirm the role of microglia during regeneration. The loss of microglia reduced early post-injury tnf*α* expression but caused accumulation of neutrophils that expressed tnf*α* later after brain lesioning to prolong inflammation hindering inflammation resolution. Overexpression of pstat3 and *β*-catenin rescued the microglial defect after injury, confirming that microglia are upstream of these signaling pathways. These findings show the critical role of microglia in regulating inflammation resolution and repairing the zebrafish brain and suggest pathways for manipulation to improve mammalian brain regeneration.

## Results

### Liposomal clodronate attenuates the microglial response to brain injury

Brain injury induces microglial activation and inflammation in both the zebrafish and mammalian CNS. Acute inflammation is largely detrimental to regeneration in mammals, while necessary in zebrafish (Ekdahl et al., 2003; Kyritsis et al., 2012; Ransohoff & Brown, 2012). We sought to directly test this idea by suppressing activated microglia at the time of brain injury in adult zebrafish. We generated a stab wound in the right telencephalon and analyzed cellular responses at different times after injury (Fig 1a, b). The left (contralateral) telencephalon provided an internal uninjured control, and unlesioned fish provided additional controls. We used a macrophage reporter line that expresses green fluorescent protein (GFP) under the macrophage expressed gene 1 promoter *tg*(*mpeg1*:*GFP*) and 4C4 immunostaining to label microglia (mpeg+, 4C4+) and macrophages (mpeg+, 4C4-) after injury. We found that both microglia/macrophages began to accumulate a few hours post lesion (hpl), the macrophages peaked at 1 day post lesion (dpl) and the macrophages numbers declined sharply at 2 dpl but remained elevated compared to the control through 4 dpl (Figs. 1c, 1d and Extended Data Fig. 1a). In contrast, ipsilesional 4C4/mpeg reporter double-labeled microglia began to accumulate at 1 dpl, the numbers peaked at 2 dpl coinciding with regeneration phase after injury, and then declined but remained slightly higher than in the uninjured brain at 3-4 dpl (Figs. 1c, 1d and Extended Data Fig. 1a). The percentage of *mpeg*-*GFP*+ cells that were co-labeled by the 4C4 antibody also peaked at 2 dpl (Extended Data Fig. 1d). Less marked increases in *mpeg*-*GFP*+, but not double-labeled, cells with a similar timecourse were seen contralaterally compared to uninjured controls (Figs. 1c and Extended Data Fig. 1a, c). We next injected the phagocytic cell toxin liposomal clodronate at the time of brain lesioning to attenuate the microglial response. Liposomal clodronate eliminated 80% of the activated microglia compared to control (PBS-containing) liposomes by 1 dpl, and the significant decrease persisted at 2 dpl in the ipsilesional hemisphere (Figs. 1e-f and Extended Data Fig. 1b). We also found significantly reduced reporter labeling of *mpeg1*:*GFP* fish at 2 dpl (Extended Data Fig. 1e). The contralateral hemisphere did not show significant differences (Extended Data Fig. 1f and g). Terminal deoxynucleotidyl transferase dUTP nick-end labeling (TUNEL) combined with 4C4 immunostaining showed more microglial cell death after liposomal clodronate injection compared to control liposomes (Figs. 1g, Extended Data Fig. 1h), confirming that clodronate eliminates activated microglia efficiently in the first 2 dpl.

**Figure 1.**
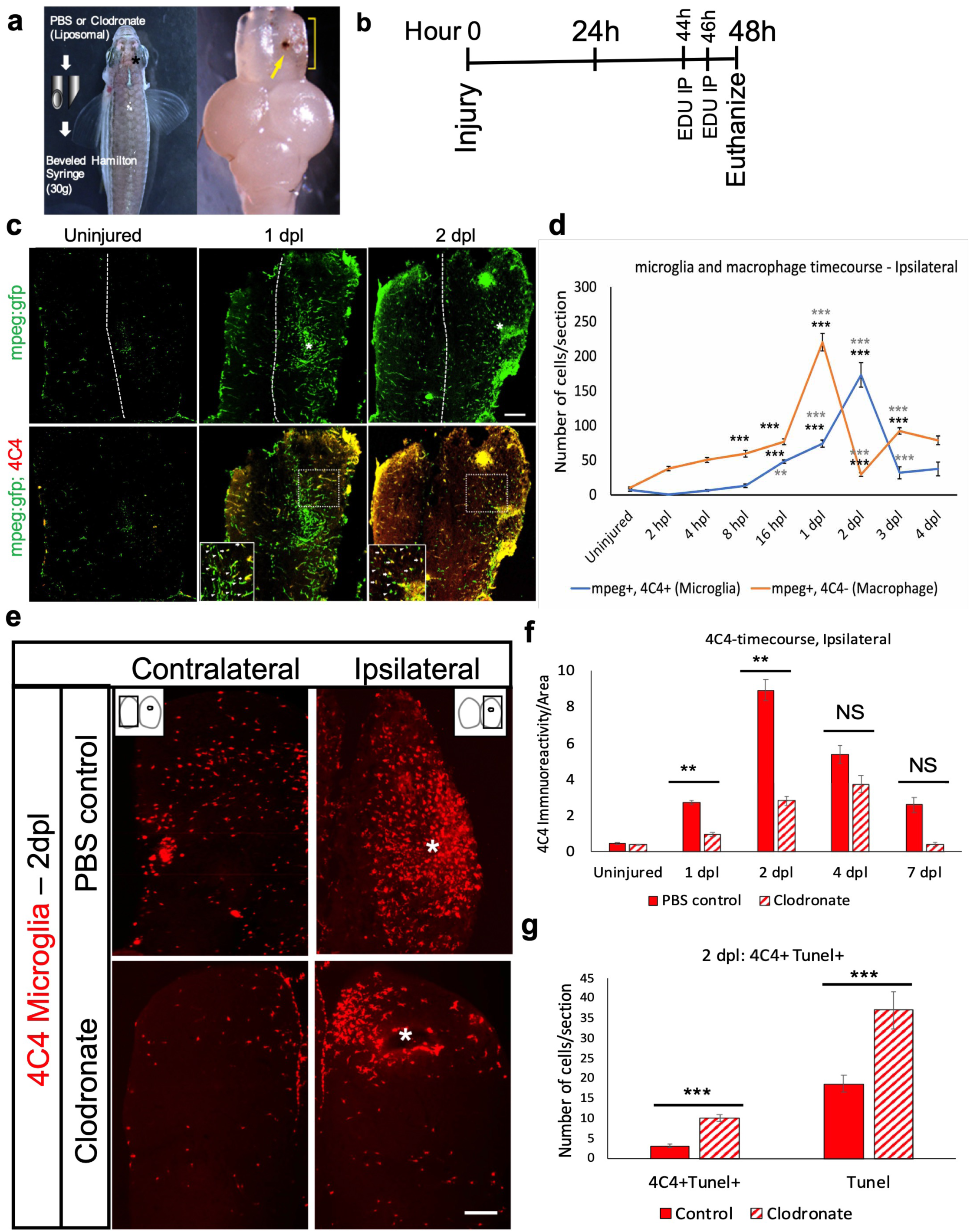
Liposomal clodronate suppresses activated microglia after telencephalic injury. a. Description of the lesioning method and injury location in the right telencephalon (asterisk, left panel), and an example of a dissected zebrafish brain showing the injury (yellow arrow, right panel). b. Timeline of lesioning, EdU labeling and euthanasia for the 2-day time point. c. Microglial response after injury at 1 and 2 days post lesion (dpl) as measured by *tg* (*mpeg*:GFP) reporter expression alone (green, top panels) or after double-labeling with 4C4 antibody to label activated microglia (red, bottom panels). The midline is indicated by the dotted lines and injured regions by asterisks in the top panels. d. Quantification of microglia represented by blue line (*mpeg:*GFP+, 4C4+ double-positive) or macrophage represented by redline (*mpeg:*GFP+, 4C4-) from 2 hours post lesioning (hpl) to 4 dpl. ***, p<0.0001 by repeated measures ANOVA. Comparisons for each curve were made between non-lesioned animals and lesioned fish at each timepoint with statistical significance denoted by black asterisks, and between lesioned fish at a given timepoint versus the previous timepoint, with dark grey asterisks denoting significant differences. e. 4C4 immunolabeling at 2 dpl after injection of control (top panels) or clodronate (bottom panels) liposomes. * indicates the site of injury. f. Quantification of 4C4+ microglia in the ipsilesional telencephalon at 1, 2, 4 and 7 dpl in control and clodronate groups. n=12, **, p<0.001 by repeated measure ANOVA. g. Quantification of 4C4 and TUNEL labeling at 2 dpl in control and clodronate groups. ***, p<0.0001 by t-test. Scale bar = 10 µm for c and e, and the error bars indicate SEM for all graphs.

### Microglial ablation inhibits injury-induced brain regeneration

We next investigated the consequences of microglial ablation on brain repair by examining brain regeneration at a later post-injury timepoint. Brains injected with clodronate or control liposomes were labeled with a nuclear stain (Bisbenzimide) at 90 dpl to assess brain structure. Remarkably, microglial ablation at the time of injury was sufficient to prevent brain repair as evidenced by the persistence of a lesion at 90 dpl in two-thirds of clodronate-treated fish, while control fish failed to show any signs of injury at 90 days and the quantification of the lesion size at different time points after injury also confirms that microglial ablation prevents brain repair (Figs. 2a-h and S2a). We also examined whether the lack of repair is associated with a glial scar that resembles that seen with mammalian brain injury. The presence of glial fibrillary acidic protein (GFAP)-expressing astrocytes and chondroitin sulfate proteoglycan (CSPG) deposition at injury sites are hallmarks of a mammalian glial scar. Interestingly, we did not observe either of these characteristics in *gfap*:GFP reporter fish (Figs. 2i and S2b) or after CSPG immunolabeling (data not shown) in the brains from fish at 90 dpl. Furthermore, the prolonged astrocytic hypertrophy (swollen processes) seen early after injury (until day 7; data not shown) appears to have resolved at later time points (Fig. 2j). These data corroborate previous findings that the zebrafish telencephalic parenchyma is devoid of canonical astrocytes and the glial scar (Silver & Miller, 2004). Taken together, our findings suggest that microglia are necessary for successful regeneration, and the lack of regeneration in the absence of microglia does not induce a glial scar.

**Figure 2.**
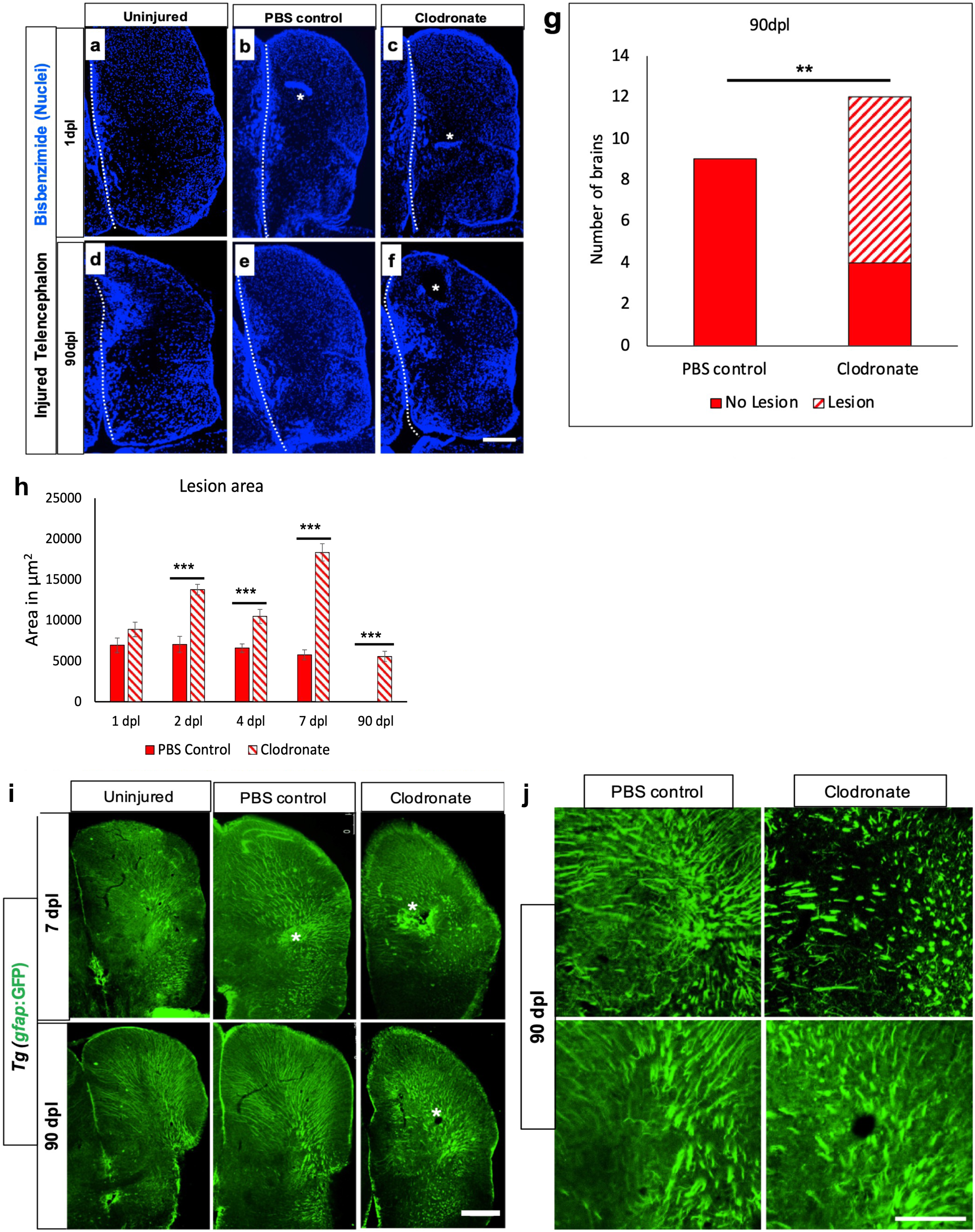
Microglial ablation inhibits brain regeneration. a-f. Nuclear staining with bisbenzimide in the right telencephalon at 1 (a-c) or 90 (d-f) dpl of uninjured (a, d), control lesioned (b, e) or clodronate-lesioned (c, f) fish. Asterisks in b, c and f denote acute (b, c) or residual (f) damage. The dashed line in each panel denotes the midline. g. Comparisons of the proportion of brains with or without residual damage at 90 dpl in control or liposomal clodronate groups. **, p = 0.0031 by Chi-Square test. h. Graph displays the lesion size quantification of control and clodronate brain at different time points. ***, p<0.0001 by repeated measures ANOVA. i. *Tg*(*gfap*:GFP) staining to assess astrocytic hypertrophy or scar- like structures at 7 and 90 dpl. Asterisks show the lesioned area/residual damage. j. Higher magnification views of *gfap*-driven reporter expression in additional fish from the 90 dpl injured control and clodronate groups. Scale bars = 100 µm.

### Microglia are essential for injury-induced adult neurogenesis

Because microglial ablation impairs brain regeneration, we next sought to determine whether the lack of repair results from a decrease in adult neurogenesis. We performed pulse-chase labeling with EdU on day 2 after injury and identified new neurons by double labeling for EdU and the neuronal marker HuC/D at 14 or 21 dpl. We found a 50% decrease in the number of EdU and HuC/D co-labeled cells at 14 and 21 dpl in brains exposed to clodronate compared to controls (Fig. 3a-d). The decrease in neurogenesis was associated with a similar percentage decrease in overall cell proliferation (Figs. 3d and S2c), indicating that neuronal differentiation was not impaired. The contralateral telencephalon did not show any significant differences at either time point (Fig. S2d, e). Thus, our data indicate that suppressing microglia attenuates injury-induced adult neurogenesis, which likely contributes to impaired brain regeneration.

**Figure 3.**
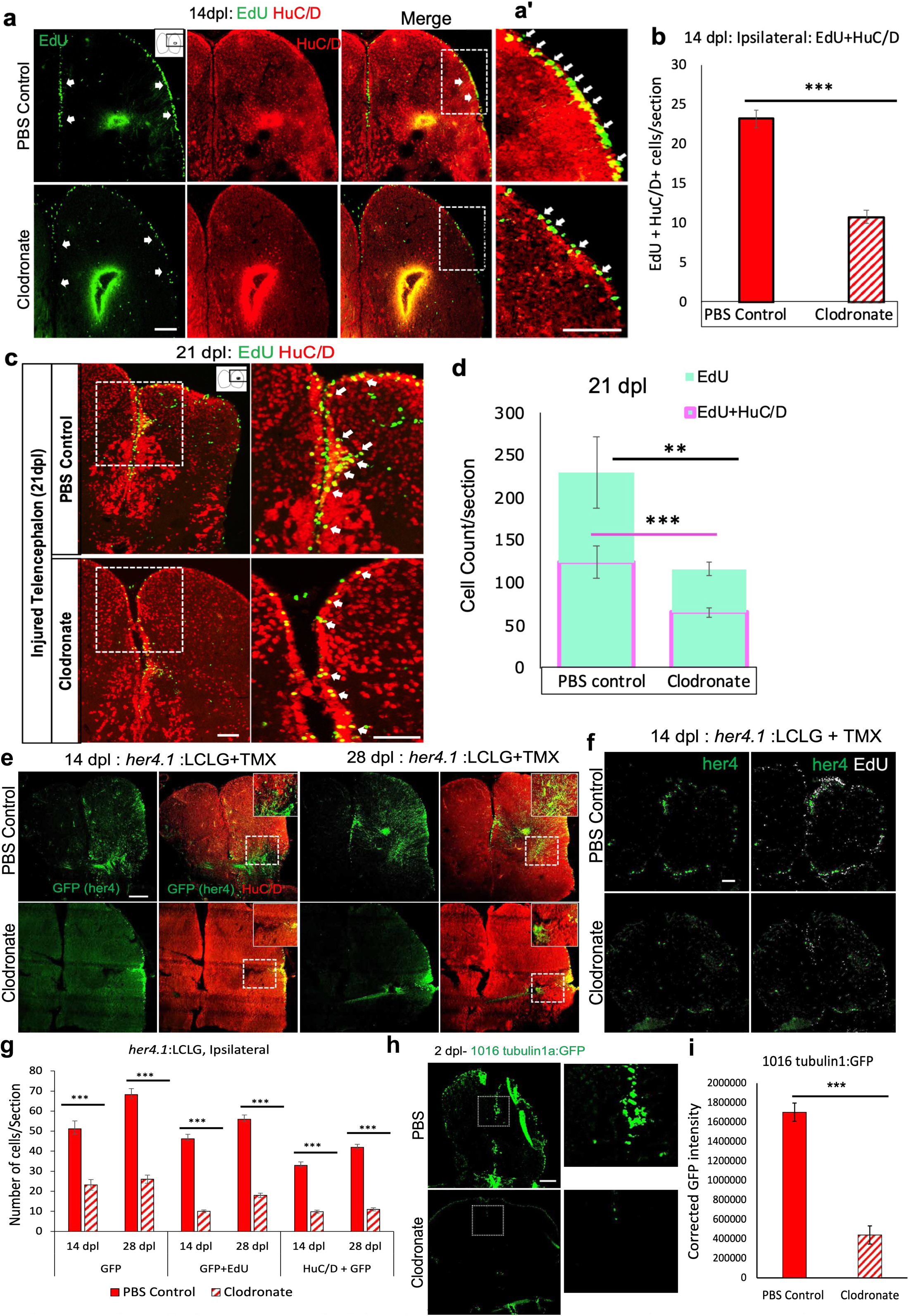
Microglial suppression impairs injury-induced adult neurogenesis. a. Confocal images of EdU staining, HuC/D immunolabeling and double labeling for EdU and HuC/D in the injured telencephala at 14 dpl after EdU administration at 2 dpl from control (top row) and clodronate injected (bottom row) brains. The arrows denote EdU+ cells in VZ regions. a’. Higher magnification images of the boxed regions in a. Note more double-labeled cells (arrows) are present in the control (top) than the clodronate-treated fish (bottom). b. Quantification of EdU and HuC/D double labeling at 14 dpl. ***, p<0.0001 by t-test. c. Confocal images of EdU and HuC/D double labeling in telencephala at 21 dpl (EdU was given at 2 dpl). Right panels show higher magnification views of the boxed regions and the arrows indicate double labeled cells. d. Quantification of EdU+ cells (green) and cells co-labeled for EdU and HuC/D (magenta outline) in injured telencephala at 21 dpl (EdU was given at 2 dpl) shows proportionally fewer cells in the clodronate-treated group. **, p<0.001 for EdU only; and ***, p<0.0001 for double-labeled cells. e. Confocal images of her4.1(GFP) and HuC/D labeling of telencephala at 14 dpl (left 4 panels) or 28 dpl (right 4 panels) from fish treated with control (top row) or clodronate (bottom row) liposomes show decreased GFP labeling and double-labeled cells in the clodronate group. Insets at upper right in some panels show higher magnification views of the boxed areas. f. Confocal images of telencephala show decreased labeling for her4.1-GFP and EdU in the clodronate-treated fish (bottom row) at 14 dpl. 4-OH-TMX was given at the time of lesioning for both f and g, and EdU was administered at 2 dpl. g. Quantification of her4.1-GFP, her4.1-GFP/EdU co-labeling and her4.1-GFP/HuC/D co-labeling at 14 and 28 dpl. *** p<0.0001. h. GFP-labeled images of telencephala from a 1016tubulin1a-GFP reporter line at 2 dpl after injection of control or clodronate liposomes. Note the marked reduction in GFP in the clodronate-treated fish. The boxed areas are shown at higher magnification in the right panels. i. Quantification of relative corrected total cell fluorescence (CTCF) intensity of 1016tubulin1a-GFP reporter expression. ***, p<0.0001. Scale bars = 100 µm for all image panels, and the error bars indicate SEM.

Adult-generated neurons in the zebrafish telencephalon arise from radial glial stem cells in the ventricular zone (VZ) that are regulated in part by Notch signaling (Kroehne et al., 2011; Than-Trong & Bally-Cuif, 2015). *her4*.1 is a Notch target gene selectively expressed by zebrafish radial glia (Kroehne et al., 2011; Skaggs et al., 2014). To lineage trace radial glial stem cell progeny, we used a double transgenic line consisting of the *her4.1* promoter driving tamoxifen-inducible Cre recombinase (*her4*.1:CreER^T2^) bred to a pan-neuronal Cre reporter line expressing GFP upon Cre mediated recombination (ß-actin2:loxp-mCherry-loxp-GFP), referred to as (ß-actin2:LCLG) (Ramachandran et al., 2010). We induced recombination by injecting 4-hydroxy-tamoxifen (4- OH-TMX) both IP and directly into the brain during the injury to increase recombination efficiency. GFP+ cells were quantified at 2, 14 or 28 dpl in both control liposome- and clodronate liposome-injected fish. We observed minimal GFP expression in the VZ on day 1, followed by markedly increased expression by 2 dpl (Fig. S2g). By day 14, GFP-positive cell numbers increased further, and the cells appeared at the injury zone to a much greater extent in controls than in the clodronate group (Fig. 3e, g). Fish not receiving 4-OH-TMX or the contralesional telencephalon after 4-OH-TMX injection did not display substantial numbers of GFP+ cells after injury (Fig. S2f, h).

To investigate whether microglial ablation affected the proliferation of *her4*-GFP lineage-traced cell progeny, we injected EdU at 2 dpl and quantified GFP/EdU double-labeled cells after 14 or 28 days. We observed that proliferating *her4*-positive cell progeny decreased significantly with clodronate treatment vs. controls by 14 and 28 dpl in the ipsilesional hemisphere (Fig. 3f and g), with no difference between groups in the contralateral hemisphere (Fig. S2h). To determine whether clodronate lesioning influenced the differentiation of *her4*+ radial glial cell progeny into neurons, we assayed GFP-positive cells for co-expression of the neuronal marker HuC/D at 14 or 28 dpl. As expected, we found that clodronate injected fish had significantly reduced GFP and HuC/D double-positive cells on the side of the injury (Fig. 3e and g), while the contralateral hemisphere showed no differences (Fig. S2h). These results further support the idea that microglia are critical for injury-induced adult neural progenitor cell proliferation and neurogenesis.

We next explored injury effect in the *Tg*(*1016tubulin1a*:*GFP*) zebrafish reporter line in which GFP is only expressed in developing or regenerating CNS neurons (Ramachandran et al., 2010; Skaggs et al., 2014). In the uninjured brain, GFP expression is minimal and localizes to a few cells in the VZ (data not shown); by day 2 after injury, however, strong VZ GFP expression is induced (Fig. 3h). We quantified GFP fluorescence intensity in clodronate-treated vs. control lesioned brains and found significantly reduced GFP expression in the clodronate group (Figs. 3h, I and S2i). This finding indicates that microglia are necessary for inducing injury responsive promoter activity like that seen in retina regeneration (Goldman & Ding, 2000; Goldman et al., 2001).

### Microglial suppression impairs injury-induced intermediate progenitor cell proliferation

The radial glial aNSCs in the VZ proliferate in response to injury. Because we found that microglial ablation impairs regeneration, we asked whether microglia are necessary for aNSC proliferation after injury. We first examined overall cell proliferation and found that microglial ablation did not change cell proliferation at 1 dpl but significantly reduced total proliferating (EdU pulse-labeled) cells in the ipsilesional hemisphere of clodronate injected brains compared to controls at 2 dpl (Fig. 4a, b). Interestingly, proliferation subsequently increased in the clodronate group at 4 and 7 dpl (Fig. 4b), suggesting that suppressing microglia early after brain injury delays the peak of the injury-induced proliferative response. No significant differences between groups were seen contralaterally at any timepoint (Fig. S3a, b). Further analyses revealed that the reduced proliferation in the clodronate group at 2 dpl was not seen in the VZ but was limited to the SVZ plus parenchymal regions (Fig. S3c, d). In contrast, the later increase in cell proliferation at 4 and 7 dpl in the clodronate injected brains was seen mainly in the SVZ plus parenchymal regions on day 4, and in all 3 areas on day 7 (Fig. S3c, d).

**Figure 4.**
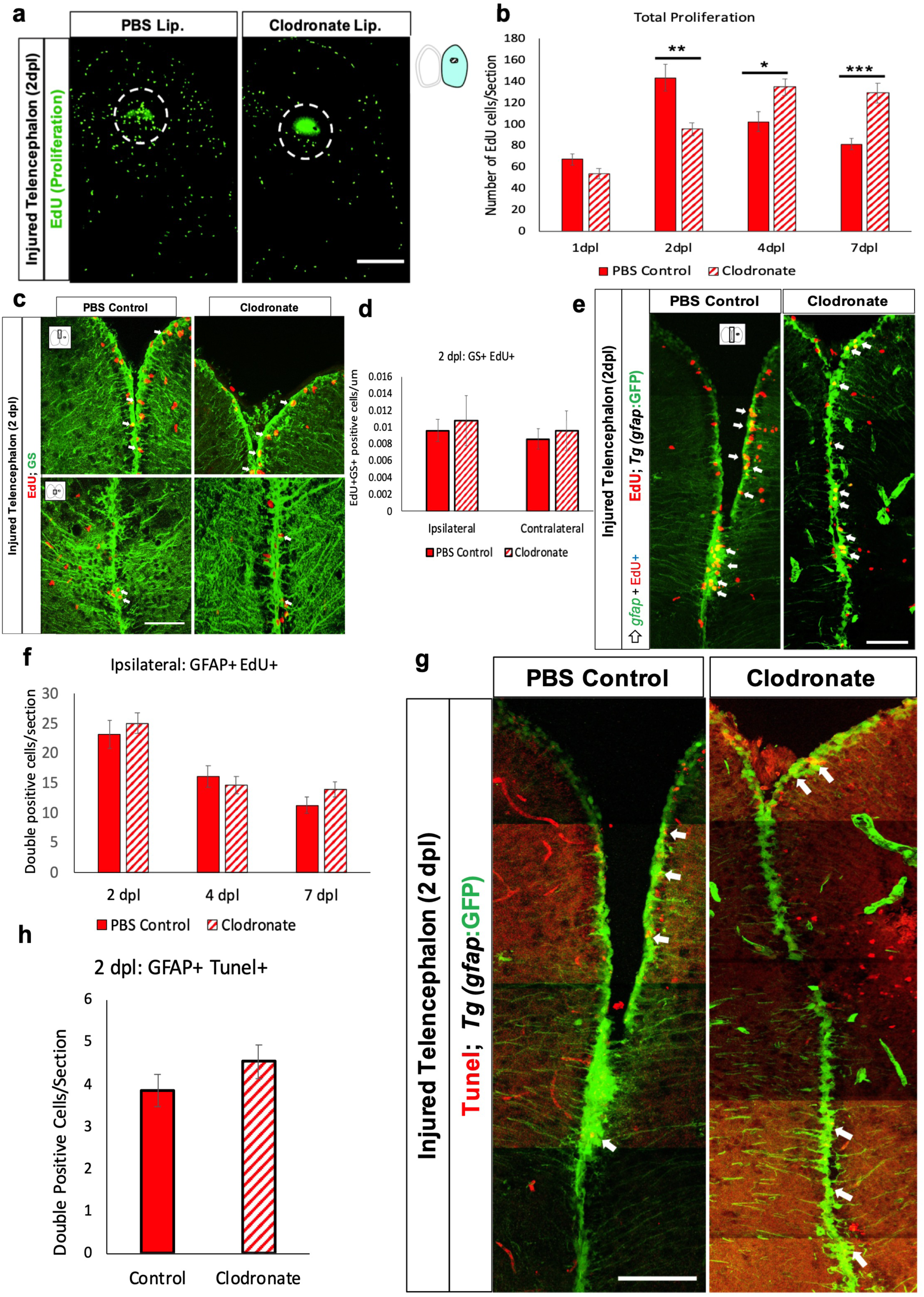
Microglia are dispensable for adult neural stem cell proliferation. a. EdU labeling of the ipsilesional telencephalon of control or clodronate injected brains at 2 dpl. The dashed circle marks the injury site. b. EdU quantification in the ipsilesional telencephala at 1, 2, 4- and 7-day post injury. n=8/group/timepoint, ***, p<0.0001; **, p<0.001; *, p<0.1, repeated measures ANOVA. c. Confocal images of Glutamine synthetase (GS) and EdU co-immunolabeling at 2 dpl showing anterior (top row) and more posterior (bottom row) telencephalic regions. The small cartoon insets show the image location in the telencephalon. d. Quantification of double labeling for GS and EdU in both ipsilesional and contralateral telencephalic hemispheres. e. Confocal images of *gfap*-driven GFP reporter and EdU co-labeling of midline telencephala at 2 dpl from clodronate and control liposome-treated fish. Arrows indicate double labeled cells. f. Quantification of ipsilesional Gfap/EdU co-labeled cells at 2, 4 and 7 dpl at the ventricular zone revealed no differences between groups. g. Confocal images of telencephalic sections co-labeled for *gfap*-driven GFP reporter and TUNEL stain at 2 dpl at the midline VZ. h. Quantification of VZ *gfap*-driven GFP reporter/TUNEL double-positive cells at 2 dpl showed no significant difference between groups. Scale bars = 100 µm in all image panels, and the error bars indicate SEM.

To specifically investigate if injury-induced aNSC proliferation is affected, we quantified EdU pulse-labeled cells that co-expressed the aNSC markers GFAP or glutamine synthetase (GS). We first examined EdU and GS co-expression and surprisingly found that clodronate lesioning did not significantly alter aNSC proliferation at 2 dpl (Fig. 4c, d). Similarly, no difference in proliferating aNSCs labeled with the *Tg*(*gfap*:GFP) reporter line were present at 2 dpl (Fig. 4e, f). We also found no significant increase in EdU/GFAP double-labeled cells vs. controls at 4 and 7 dpl (Fig. 4f). Given the lack of changes in aNSC proliferation at 2 dpl, it is unlikely that clodronate induced aNSC death. Consistent with this idea, no significant difference arose in VZ GFAP/TUNEL double-labeled cells between control and clodronate groups (Fig. 4g, h). Moreover, fluorescently labeled control liposome injection during injury revealed that the liposomes minimally cross the midline to the contralateral side, and they appeared to be only rarely taken up by aNSCs (Fig. S3g, h). As predicted, we observed that most GFP-positive microglia were labeled with or adjacent to the fluorescent liposomes, consistent with the idea that clodronate liposomes are specifically taken up by phagocytic microglia/macrophages (Fig. S3i). Together, these findings indicate that microglial suppression with clodronate does not attenuate aNSC proliferation and suggest that clodronate does not induce aNSC death.

Given the lack of changes in aNSCs after clodronate treatment, we next examined proliferation rates of GFAP-negative progenitors in the VZ/SVZ. We found a 30% decrease in non-GFAP-expressing dividing cells (GFAP^-^/EdU^+^) in the VZ/SVZ of clodronate-injected brains (Figs. 5a, b). We asked whether the GFAP-negative dividing cells that are decreased by microglial ablation comprise the Sox2-positive amplifying intermediate progenitor cell (aIPC) population. We quantified Sox2/EdU double-labeled cells at 2 or 4 dpl and found that the clodronate-treated group had a greater than 50% reduction in proliferating Sox2+ cells (Figs. 5c, d), likely accounting for the reduction in GFAP-negative cell proliferation. Furthermore, Sox2+ aIPC proliferation remained suppressed at 4 and 7 dpl (Fig. 5d, S3e), and the total number of Sox2+ cells was significantly decreased in the clodronate group at both 2 and 4 dpl (Fig. 5e, f). To explore whether the reduction of total Sox2-labeled aIPCs reflected progenitor cell death, we used TUNEL staining and found no significant difference in the small numbers of Sox2/TUNEL double-positive cells between control and clodronate groups (Figs. 5g, S3f). Together, these findings indicate that microglial ablation reduces the post-injury aIPC population through reduced proliferation of SOX2+ aIPCs.

**Figure 5.**
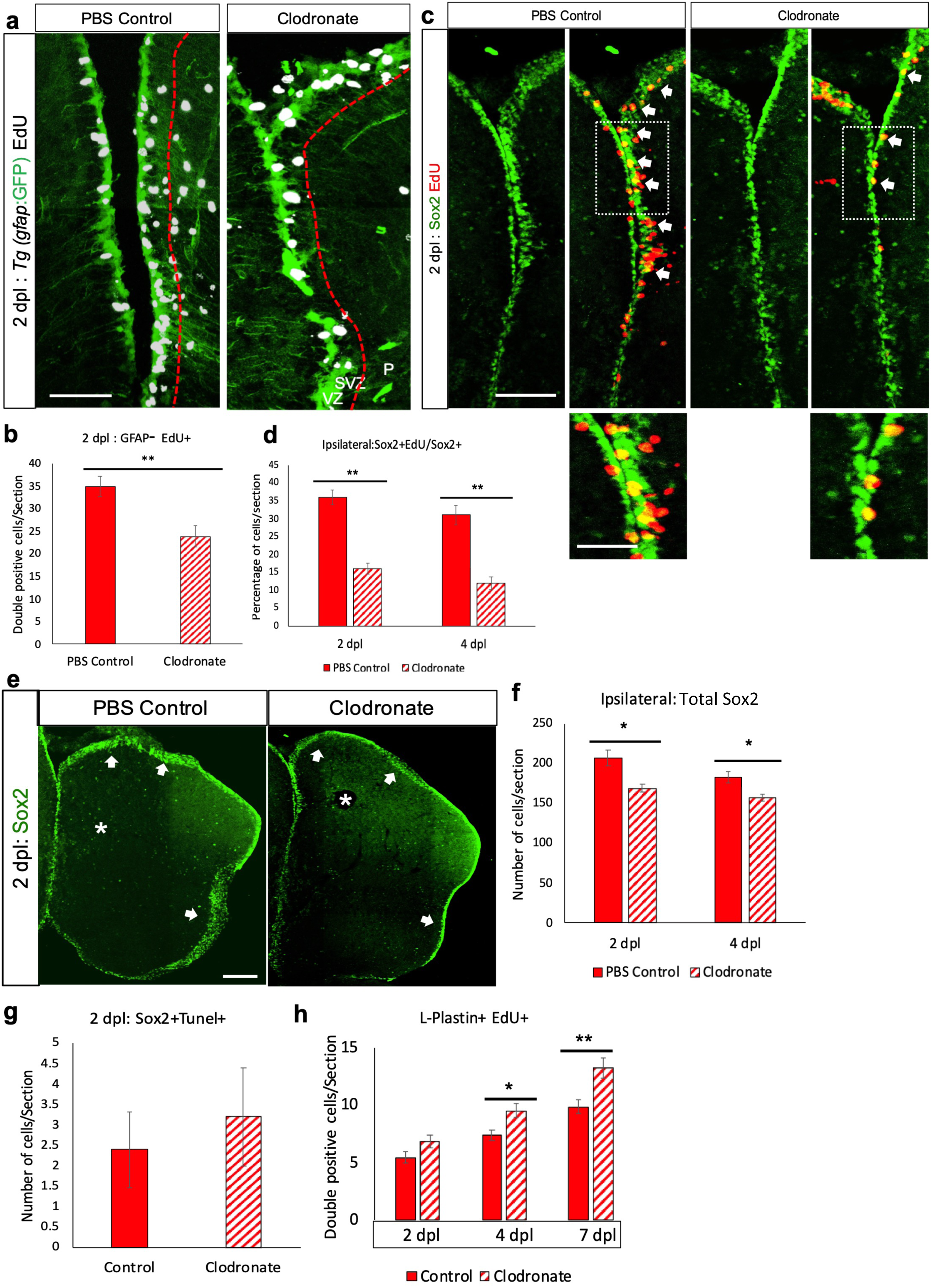
Microglial ablation impairs injury-induced intermediate progenitor cell (IPC) amplification. a. Confocal images of midline telencephalic sections from clodronate and control liposome injected fish double labeled for a *gfap*-driven GFP reporter (green) and EdU (white) at 2 dpl. The dashed red lines divide the VZ + SVZ (left of the lines) from the parenchyma (P; right). b. Quantification of *gfap*-driven GFP reporter-negative/EdU-positive cells at 2 dpl in the VZ plus SVZ. **, p<0.001 by t-test. c. Confocal images of double immunolabeling for Sox2 (green) and EdU (red) at 2 dpl in the VZ+SVZ at the midline telencephalon. Arrows indicate double-positive cells, and the boxed areas are shown at higher magnification below. d. Quantification of percentage of Sox2+ cells that co-labeled with EdU in the ipsilesional telencephalon at 2 and 4 dpl. **, p<0.001 by t-test. e. Sox2 immunolabeling of lesioned control liposome and clodronate liposome treated fish telencephala at 2 dpl. Arrows indicate labeled progenitor cells in the VZ/SVZ and asterisks denote the damaged regions. f. Quantification of Sox2-labeled cells in control and clodronate injured brain at 2 and 4 dpl. *, p<0.01 by t-test. g. Quantification of Sox2/TUNEL co-labeled cells at 2 dpl in control and clodronate groups. h. Quantification of L-Plastin and EdU double-labeled cells at 2, 4 and 7 dpl in control and clodronate injected groups revealed significantly increased double-labeled cell numbers in the clodronate group at 4 and 7 dpl. **, p<0.001; *, p<0.01. Scale bar = 100 µm for all image panels except 50 µm for the higher magnification images in panel c; error bars indicate SEM.

We next explored whether clodronate treatment influenced the proliferation of non-progenitor populations in the brain parenchyma by double-labeling for EdU and other cell-type specific markers. We found no difference in the percentage of proliferating 4C4+ microglia at 2 dpl (Fig. S4a, b), albeit with fewer total numbers of activated microglia in the clodronate group (Fig. 1f). The percentage of proliferating microglia also showed no significant difference at 4 or 7 dpl (Fig. S4b). Similarly, we did not observe any significant difference in olig2-positive oligodendrocyte proliferation or cell death in clodronate-injected vs. control fish (Fig. S4c-e). Interestingly, the clodronate group displayed significantly increased proliferation of L-plastin-positive leukocytes at 4 and, to an even greater extent, 7 dpl (Fig. 5h). The increase in leukocyte proliferation on days 4 and 7 may explain the increased overall cell proliferation at these timepoints in the clodronate group (Fig. 4b), especially in the parenchymal region. Together, our findings show that microglia suppression attenuates amplifying neural progenitor cells and stimulates leukocyte proliferation after brain injury.

### Genetic mutants for *irf8* and *csf1ra&b* (*csf1rdm*) confirms impaired injury-induced regeneration

To confirm that the clodronate effects on regeneration resulted from microglial ablation, we used two different genetic mutants that are devoid of microglia. Interferon-8 (*irf8*) is essential for microglial specification and the mutants show very few microglia and more number of neutrophils (Li et al., 2011; Shiau et al., 2015),while colony stimulating factor 1 receptor (*csf1ra&b*) is necessary for microglia development, colonization, and survival (Oosterhof et al., 2018). Here we used *irf8* (st95) homozygous mutants and heterozygous mutants as control, *csf1rdm* (double mutant), which is mutant for both *csf1ra* and *csf1rb*. Both mutants showed very few microglia in uninjured brain, but the injured brain from both mutants show drastic reduction in microglia and EdU-positive proliferating cells compared to wildtype or corresponding heterozygous mutant (Fig. 6a, b, c, d and e). We further used csf1r inhibitor PLX3397 to confirm the role of microglia during regeneration. PLX3397 treatment markedly reduced 4C4- and L-Plastin-immunoreactive cells at 2 dpl compared to vehicle-treated fish (S5a, c, g, h). We next quantified the number of proliferating cells with EdU and found a significant reduction in PLX3397-treated fish (S5b, c). As expected, the overall numbers of 4C4-, L-Plastin- and EdU-labeled cells were lower in the contralateral hemispheres and showed similar percentage reductions after PLX3397 exposure (Fig. S5e, f). PLX3397 treatment also significantly reduced aIPC proliferation at 2 dpl and neurogenesis at 28 dpl (S5i, j). The contralateral hemispheres showed similar differences, albeit with lower overall numbers (Fig. S5 k, l). Thus, both mutants, clodronate and PLX3397 treatment confirms the finding that absence of microglia blocks telencephalic injury-induced cell proliferation and neurogenesis, supporting the importance of injury-activated microglia in brain reparative processes.

**Figure 6.**
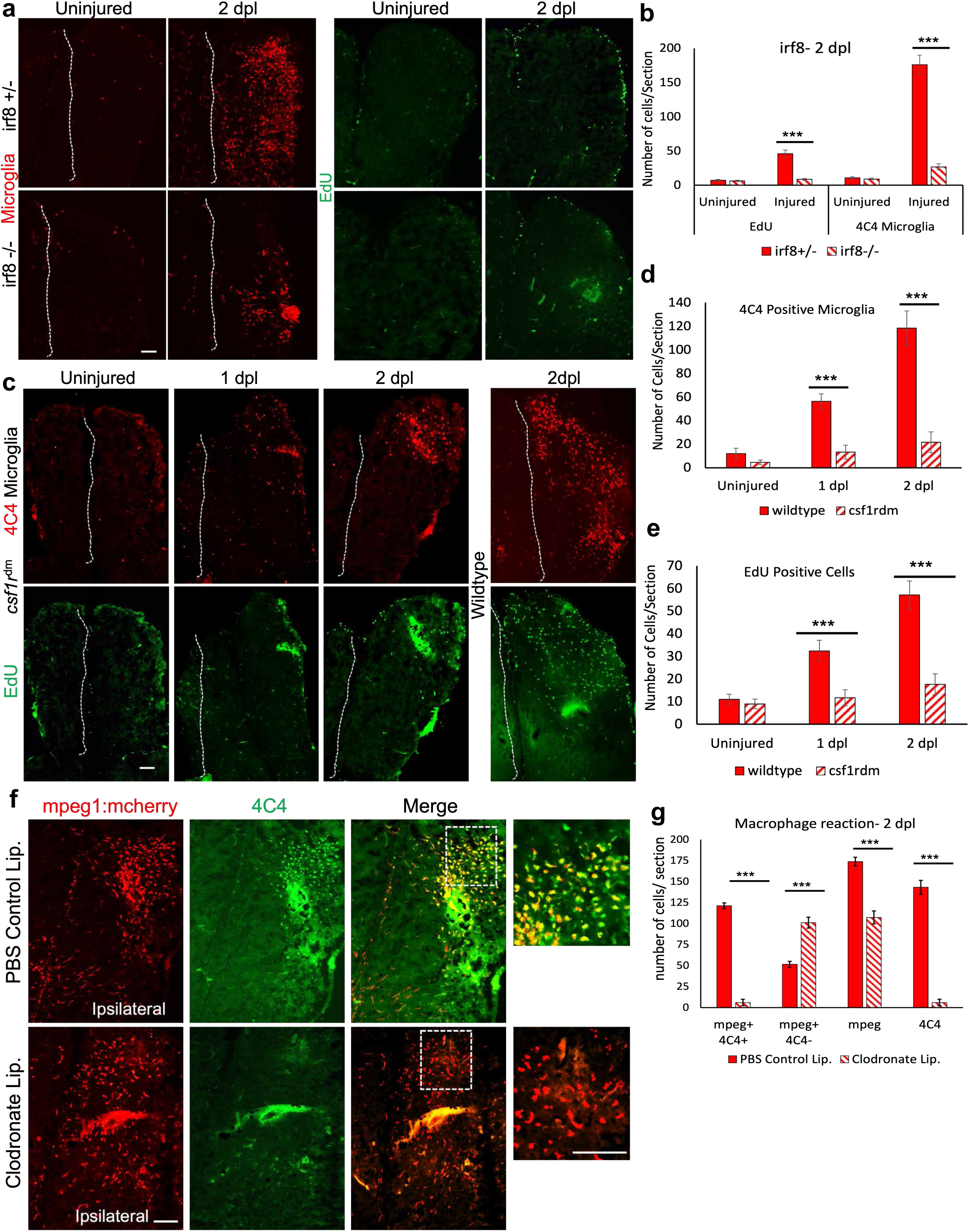
Genetic mutants for *irf8* and *csf1ra&b* (csf1rdm) display reduced microglial population, failure injury-induced cell proliferation, and macrophages are not sufficient for regeneration. a. Staining for 4C4 (microglia) and EdU labeling (proliferating cells) of *irf8* heterozygous (+/-) and homozygous (-/-) fish brain with or without injury at 2dpl. b. Quantification of microglia and EdU-positive cells in *irf8* mutants and the control at 2dpl. c. Staining for 4C4 (microglia) and EdU (proliferating cells) of *csf1rdm* mutant and wildtype brain with or without injury. Dashed lines indicate the midline in a and c. d. The graph displays the quantification of microglia in csf1rdm and wildtype brains. e. The graph displays the quantification of EdU-positive cells in *csf1rdm* and wildtype brains. f and g, shows quantification of macrophages and microglia after clodronate treatment. f. Double staining for mcherry to label mpeg1-positive cells from *mpeg1*: mcherry reporter fish and 4C4 for microglia in control and clodronate injected fish. The images are ipsilateral side and both single channels, merged view. The dotted box is shown at higher magnification. g. Quantification of macrophage (mpeg+, 4C4-) and microglia (meg+, 4C4+) in control and clodronate brain at 2 dpl. ***, p<0.0001. Scale bars = 100 µm for all image panels, and the error bars indicate SEM.

### Macrophages are not sufficient to induce brain regeneration without microglia

To address if macrophages are necessary and sufficient for injury-induced brain regeneration and if clodronate treatment affects invading macrophage population after brain injury, we have used clodronate treated brain where activated microglia are ablated or any phagocytic cells around the injury zone and a macrophage reporter line *mfap4*: dLanYFP-CAAX (Walton et al., 2015). Macrophages are peripheral immune cells that invade CNS after blood-brain barrier breakage after brain injury. Macrophages are shown necessary for regeneration of spinal cord and heart and heart in zebrafish while microglia (4C4 positive) are not (Lai et al., 2017; Themistoklis M. Tsarouchas et al., 2018). We notice that macrophages enter the CNS few hours after brain injury, peaking at 1dpl, but their numbers are sharped reduced at 2 dpl, when the regeneration phase is at peak (Fig. 1c, d). We found that the clodronate treated brain only shows drastic reduction in microglia (mpeg+, 4C4+), while macrophages (mpeg+, 4C4-) still present at significant numbers in the clodronate brain, indicating that the macrophages might not be enough to induce regeneration after injury (Fig. 6f, g). Further, we also learned that mfap4 reporter line expression does not show any significant reduction in macrophage quantity after clodronate treatment in CNS (S5m, n). This result further confirms that the activated microglia are necessary for brin regeneration while macrophages are not sufficient.

### Suppressing microglia after brain injury attenuates pro-regenerative signaling

Inflammatory cytokines and signaling molecules such as *stat3*, *ascl1a*, *lin28* and ß -*catenin* are known to play a crucial role during regeneration of zebrafish fin, heart and retina (Becker & Becker, 2015; Elsaeidi et al., 2018; Yi Fang et al., 2013; Lotan & Schwartz, 1994; Wan et al., 2014; Wehner et al., 2014; Zhao et al., 2014). Hence, we examined whether microglia are necessary to activate these pro-regenerative signaling pathways to promote repair. Gene expression analysis via qRT-PCR showed that clodronate-treated brains had reduced expression of the inflammatory *cytokines tnfa, il1ß, il11a, il12* and *arginase1*, and the signaling molecules *stat3, ascl1a, lin28 and ß -catenin* were also reduced at 1dpl (Fig. 7a). Timecourse analysis of gene expression via qRT-PCR for *stat3*, *ascl1a* and c-*myc* (b-catenin target) and *gata3,* a transcription factor induced after injury *(Kizil, Kyritsis, et al., 2012)* in clodronate, and the two genetic mutants showed significantly reduced expression at multiple time points compared to the control or heterozygous mutants (Fig. 7d-g). We validated the altered expression for a subset with reporter lines for phosphorylated pStat3 [*Tg(gfap: stat3*-GFP)], *ascl1a* [*Tg(ascl1a:GFP*)] and Wnt/ß-Catenin signaling [*Tg(tcf7miniP:2dGFP*)], and confirmed that microglial ablation strongly reduces Stat3 and *ß*-Catenin signaling, while *ascl1a* was more modestly decreased (Figs. 7b, c and S6a). Moreover, the radial glial reporter line *Tg*(*gfap*:GFP) showed slight change after clodronate injection (Figs. 7c and S6a), consistent with other results (Fig. 4e, f). These findings show that microglia are critical upstream activators of these pro-regenerative signaling pathways after injury.

**Figure 7.**
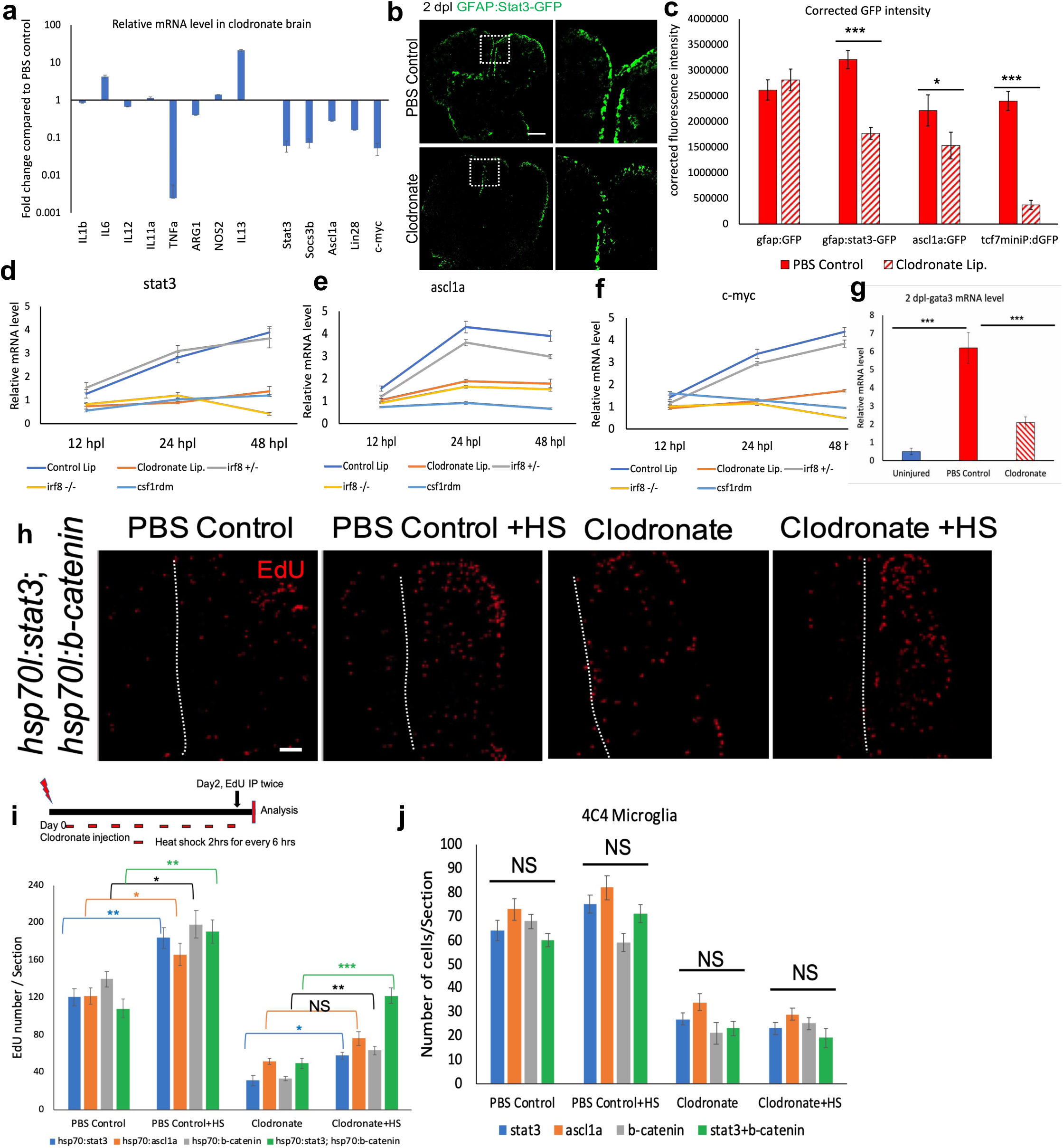
Microglial ablation after brain injury attenuates pro-regenerative signaling. a. qRT-PCR showing relative mRNA levels of different inflammatory cytokines or regenerative signaling molecules. Two pooled telencephala (injured hemispheres only) per condition (clodronate-treated vs. control lesioned) were used, and the Y axis is log fold change. Error bars indicate SD. b. GFP labeling of telencephala from a pStat3 signaling reporter line at 2 dpl after control or clodronate liposome injections. The boxed areas in the left panels are shown at higher magnification at right. c. Quantification of corrected total cell GFP immunolabelling intensity of right (injured) telencephala from *tg*(stat3-GFP), tg (gfap:GFP), tg(ascl1a:GFP) and tg(tcf7miniP:dGFP) reporter fish at 2 dpl after clodronate injury compared to controls. d-f. qRT-PCR timecourse analysis of gene expression for *stat3*, *ascl1a*, *c-myc* in clodronate treated, *irf8* and *csf1rdm* mutant brains. g. qRT-PCR for *gata3* at 2dpl after clodronate. f-j. Rescue analysis after clodronate injury with ectopic overexpression of pro-regenerative proteins. h. EdU labeling of control and clodronate injected brains with or without heat shock (HS) to induce stat3 and ß-catenin transgene expression. i. Quantification of EdU+ cells at 2 dpl after ectopic activation of stat3, ascl1a, ß-catenin and stat3+ ß-catenin with or without HS treatment in control and clodronate injured fish. The schematic above shows the experimental design. j. Quantification of 4C4-immunolabeled cells at 2 dpl after injury with or without HS in the clodronate rescue experiments. NS, not significant; ***, p<0.0001; **, p<0,001; *, p<0.01 by t-test in c, d, f and g. Scale bars = 100 µm for image panels, and error bars indicate SEM and SD in a, d, e, f and g.

### Stat3 and ß-Catenin pathway activation rescues the microglial ablation proliferation defect

Given our results that microglia are upstream of these signaling pathways, we sought to determine whether ectopic activation of Stat3, Ascl1a and ß-Catenin signaling alone or in combination would rescue the proliferation defect seen after microglial ablation. We overexpressed constitutively-activate (CA) Stat3 [Tg*(hsp70l:CA-stat3*)], ascl1a [Tg*(hsp70l:ascl1a*)] and ß-catenin [Tg*(hsp70l:CA-ß-catenin*)] with multiple heat-shocks for two days after injury in control and clodronate liposome-injected fish. Additional control and clodronate injury groups without heat shock were also examined. Individually, stat3 and ß-catenin, but not ascl1a, overexpression showed a modest but significant increase in proliferation at 2 dpl in heat shock-treated clodronate-injured brains compared to those without heat shock (Fig. 7i, S6b). We next used a double Tg line [Tg*(hsp70l:CA-stat3; hsp70l:CA-ß-catenin*)] to ask whether combined stat3 and ß-catenin overexpression would reverse the clodronate-induced proliferation defect after injury. Indeed, we found a more marked increase in proliferation in the double transgenic line with heat-shock compared to no heat-shock, restoring proliferation to control levels (Figs. 7h, i and S6b). In addition, *stat3* or *ß–catenin* overexpression did not influence microglial numbers in control- and clodronate-lesioned fish (Fig. 7j), and heat-shock alone did not induce proliferation and did not alter microglia numbers in wildtype with or without injury (Fig. S6c). We further confirmed that all the over expression lines indeed express the intended genes after heat shock via qRT-PCR (S6d). Together, these results show that Stat3 and ß–Catenin signaling are activated by microglia and in combination are sufficient to rescue the proliferation defect caused by microglial ablation.

### Microglial-dependent *tnf⍺* expression is necessary to activate Stat3 and ß-Catenin signaling

Our gene expression analysis revealed that *tnfα* and *il1b* mRNA is prominently downregulated after lesioning plus microglial ablation (Fig. 7a). To determine whether the microglial-stimulated inflammatory response after injury links to regeneration through tnf*α*, we investigated changes in *tnfα* expression patterns after injury. Timecourse analysis of *tnfα* and *il1b* gene expression in clodronate brain and in the genetic mutants without microglia showed significantly reduced expression after injury, specifically *tnfα* (Fig. 8a, b . *tnfα* expression is induced within an hour after injury and remains strongly expressed for 12 hours (about 14-fold over control) before starting to decline at 24 hours, in contrast, *tnfa* expression were reduced early on without microglia but the expression was increased significantly at 2 dpl on wards compared to control and the genetic mutants display greater increase then the clodronate brain (Fig. 8a).

**Figure 8.**
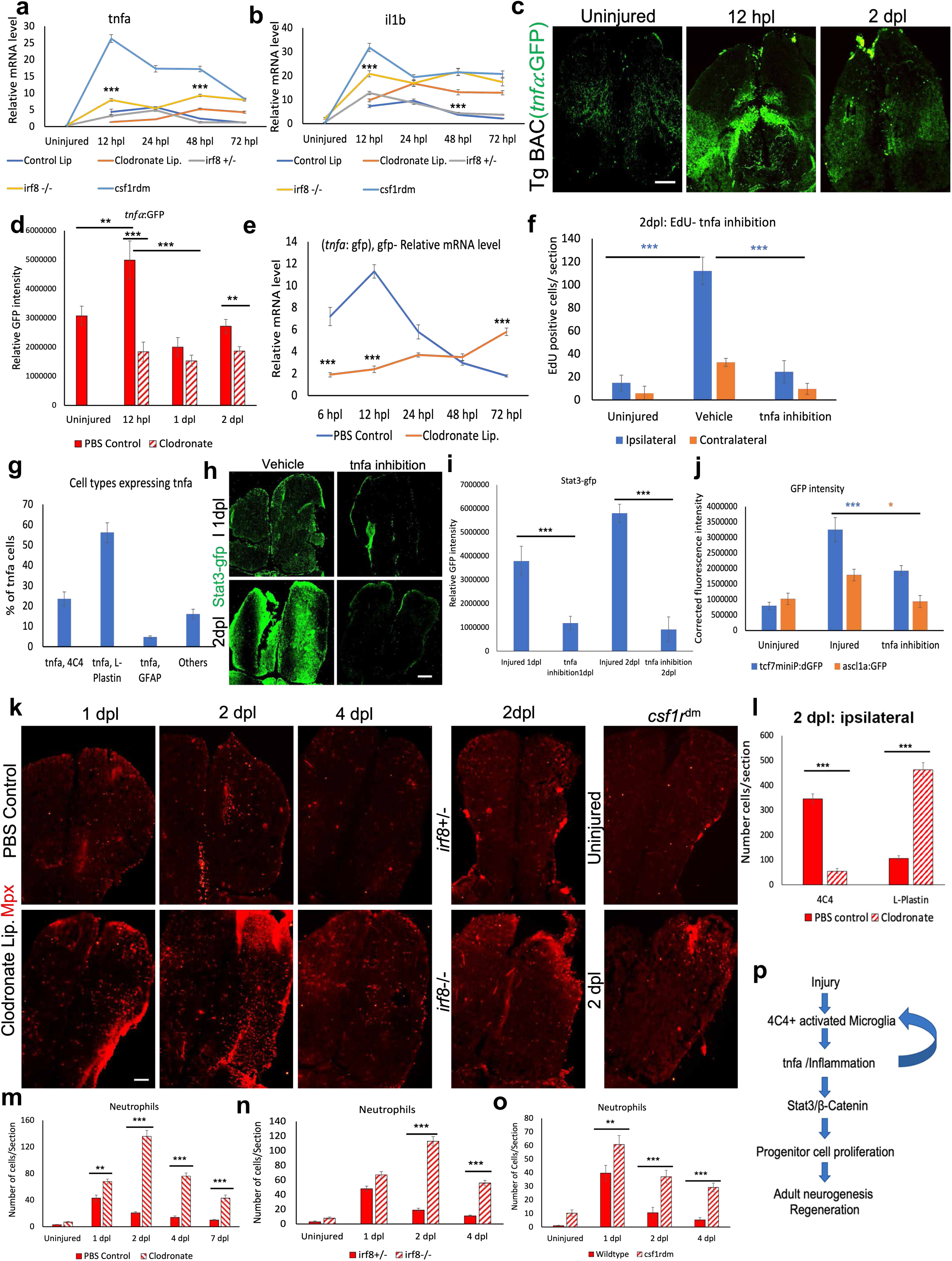
Microglial-dependent tnfa expression is necessary to activate stat3 and ß-catenin signaling to induce regeneration. a-b, qRT-PCR timecourse analysis showing relative mRNA levels of *tnfa* and *il1b* in control, clodronate, *irf8* and *csf1rdm* mutant brains after injury. Five telencephala (only right hemisphere) pooled from injured or uninjured brain were used. Error bars indicate SD. Stars indicates the statistical difference between control and clodronate or control and the mutant at given time point. Clodronate and *csf1rdm* compared to control, and *irf8*+/- compared to *irf8*-/-. c. Confocal images of GFP reporter labeling *Tg*BAC(*tnfa*:GFP) for *tnfa* expression in an uninjured fish and at 12 hpl and 2 dpl. d. Quantification of GFP corrected intensity from *tnfa*:GFP reporter line at 12 hpl, 1 dpl, and 2 dpl in control or clodronate liposome groups. e. qRT-PCR for gfp mRNA from *tnfa*:GFP reporter fish after clodronate treatment. The stars indicate the significance between control and clodronate at given timepoint. f. Quantification of EdU+ cells after injury with or without thalidomide (tnfa inhibitor). DMSO vehicle injection was used to injure the control brain. g. Quantification of cells double labeled for GFP (*tnfa:GFP*) and other cellular markers, shown as percentage of total tnfa+ cells. h. Confocal images of *stat3*-GFP reporter expression at 1 (top row) or 2 (bottom row) dpl in groups exposed to vehicle (left panels) or the tnfa inhibitor thalidomide (right panels). i. Quantification of GFP intensity from the ipsilesional hemisphere in *stat3*:GFP reporter fish at 1 and 2 dpl with or without thalidomide. j. Quantification of GFP immunolabeling intensity from the ipsilateral side of *ascl1a*:GFP and *tcf7miniP*:dGFP transgenic reporter lines in uninjured fish or in lesioned fish with or without thalidomide. k. immunolabelling control, clodronate treated, *irf8* and *csf1rdm* brains for Mpx (neutrophils) after injury. l. Quantification of microglia and leukocytes in the ipsilesional hemi-telencephala of control or clodronate liposome injected fish at 2 dpl. m-o. Quantification of Mpx-positive neutrophils from control, clodronate treated, *irf8* and *csf1rdm* mutant brains. p. Proposed model of the microglia-induced signaling pathway during injury-induced telencephalic regeneration in zebrafish. ***, p<0.0001; **, p<0,001; *, p<0.01 by t-test for panels a, d, e, h, j and l. Scale bars = 100 µm, and the error bars indicate SEM except for panel a, b, e.

To further validate *tnfα* expression changes, we used a bacterial artificial chromosome-engineered transgenic zebrafish reporter line *Tg*BAC*(tnfa:GFP)* (Marjoram et al., 2015), hereafter referred as *tnfa*:GFP, that reports *tnfα* transcriptional activity. Expression of the *tnfα* reporter is low in the uninjured brain but markedly increases by 12 hpl, returning to basal levels by 1 dpl (Fig. 8c, d). We next examined whether microglia are necessary for the injury-induced induction of *tnfα*. Interestingly, microglial ablation significantly reduced GFP expression at 12 hpl, indicating a lack of injury-induced upregulation of tnf*α*, and slightly increased levels at 2 dpl in the clodronate group, however still lower than the control level contrast to the *tnfa* gene expression via qRT-PCR (Fig. 8c, S7b). To eliminate that the GFP protein translation and stability does not misrepresent the dynamic gene expression of *tnfa*, we quantified gfp mRNA level in *tnfa*:GFP fish after clodronate. As expected, the qRT-PCR showed that gfp mRNA expression are consistent with *tnfa* mRNA expression, which is reduced at earlier timepoints after injury without microglia but increased significantly after 2 dpl on wards (Fig. 8e). These findings indicate that early microglial activation is necessary for injury-induced *tnfα* expression and the necessity of tight regulation of inflammation. As expected, since inflammation is necessary for regeneration in fish, tnf*α* is also required for cell proliferation after injury in zebrafish, as treatment with two different tnf*α* inhibitors (thalidomide and GIT27- 4,5-Dihydro-3-phenyl-5-isoxazoleacetic acid) drastically suppressed injury-induced cell proliferation (Figs. 8f and S7c). Double labeling revealed that tnf*α* was co-expressed by L-Plastin-positive leukocytes (55% of tnf*α*+ cells) and to a lesser extent by 4C4+ microglia (26%), with very few GFAP-positive radial glia showing co-localization (3%) (Fig. 8g, and S7a), suggesting that the majority of tnf*α* in the injured brain comes from infiltrating leukocytes but microglia contributes as well, probably inducing very early after injury before the peripheral immune cells invade.

We next examined whether Tnf*α* is necessary to activate downstream signaling during regeneration. We found that injury-induced Stat3 and ß-Catenin signaling were strongly suppressed, while Aslc1a was more modestly but significantly reduced after Tnf*α* inhibition with thalidomide (Figs. 8h-j and S7f, g). Furthermore, injecting the thalidomide during injury reduced *stat3, ascl1a, lin28* and c*-m*yc (the latter a ß-catenin target gene) expression at 2 dpl compared to vehicle (Fig. S7e), further confirming that Tnf*α* is necessary to activate pro-regenerative signaling pathways. Tnf*α* inhibition did not affect the 4C4+ microglial population after injury (Fig. S7d, h), supporting the idea that microglia are upstream of Tnf*α* signaling during regeneration. Together, our results suggest that microglial-induced Tnf*α* is necessary to activate downstream pro-regenerative signaling to promote telencephalic repair.

### Microglia are crucial for inflammation resolution

So far, our results show that the microglia are necessary to induce inflammation and the downstream pro-regenerative signaling via inducing *tnfα* expression after injury. Also, we learnt that absence of microglia leads to increased and extended period of *tnfα* and *il1b* expression than the control. Given that the resolution of inflammation typically dictates the degree of repair, we next explored whether microglia are necessary to remove proinflammatory peripheral immune cells (neutrophils) to attenuate ongoing inflammation. We showed that L-plastin cells proliferate more in the clodronate brain parenchymal region compared to control (Fig. 5h). we also notice leukocytes accumulation in the clodronate brain without 4C4-positive microglia at 2dpl (Fig. 8l and S7i). Further, to understand if microglia play a role in neutrophils clearance to suppress inflammation after 1 dpl, which is one of major steps in inflammation resolution, we quantified neutrophils in clodronate brain and the two genetic mutants. Indeed, we discovered that microglial absence leads to strong accumulation of neutrophils after 1 dpl on wards (Fig. 8k-o). The accumulation of neutrophils because of failure in neutrophil clearance after pro-inflammatory phase to activate the pro-regenerative program could also explain the increase and extended duration of *tnfα* and *il1b* expression without microglia. Also indicates the requirement of dynamic and tight regulation of inflammation during regeneration. This result suggests that the presence of activated microglia is crucial to remove proinflammatory cells to resolve inflammation during regeneration. Together, our findings show that microglia are crucial for inflammation, the resolution of inflammation, and essential for aIPC amplification and activation of pro-regenerative signaling during telencephalic regeneration.

## Discussion

Given the robust regenerative capacity of the zebrafish brain, a critical issue is to understand the unique mechanisms underlying injury-induced brain repair in zebrafish. Here we focused on the effects of inflammation by attenuating the microglial response to injury. In our adult zebrafish brain injury model, we found that suppressing activated microglia impaired brain regeneration and often led to persistent lesions, comparable to the chronic damage seen after mammalian brain injury. This defective repair was associated with decreased aIPC proliferation and neurogenesis. We also found that microglia play an important role upstream of tnfa expression and pro-regenerative signaling pathways. Importantly, microglial absence prevented neutrophil clearance and inflammation resolution. Furthermore, macrophages and neutrophils are not sufficient for regeneration. Thus, our findings demonstrate the requirement of microglia for adequate brain repair through anti-inflammatory and pro-regenerative mechanisms.

### Activated microglia play an M2-type/anti-inflammatory role to promote regeneration

Resident immune cell, microglia react after injury and promotes regeneration by inducing inflammation, regulating inflammation resolution, and activating pro-regenerative pathways. M1-type microglia are considered pro-inflammatory, while M2-type are considered anti-inflammatory and neuroprotective after mammalian brain injury (Martinez & Gordon, 2014; Ransohoff, 2016). M2-type macrophages are also involved in zebrafish cardiac and fin regeneration, where they accumulate near the injury by 1 dpl and do not express Tnfα (Lai et al., 2017; L. Li et al., 2012). We find similar results in our brain injury model, in which 4C4+ microglial accumulate strongly by 1 dpl, peak at 2 dpl and few express *tnfα*, suggesting that 4C4+ microglia are pro-regenerative/M2-type microglia. The majority of *tnfα* expression instead appears to arise from peripheral immune cells. Inflammation is necessary to induce regeneration in zebrafish. We find that, microglial ablation, in irf8 and csf1rdm mutants reduced *tnfα* expression early but later stronger *tnfα* and il1b expression, indicating that activated microglia might play a dual role during the initiation and resolution of the inflammatory response. Hence dynamic and tight regulation of inflammation is crucial for successful regeneration and microglia seems leads the way.

Unlike in the mammalian brain, we found no glial scarring in the fish despite the chronic damage and failed inflammation resolution induced by microglial suppression. The lesioned zebrafish did show early radial glial hypertrophy that attenuated within two weeks. The absence of a scar may reflect the lack of parenchymal astrocytes in zebrafish, or some innate system to overcome scarring despite the presence of a persistent lesion, even though astrocyte-like cells were reported recently in developing zebrafish brain but yet to be found in adult telencephalon (Chen et al., 2020). Thus, suggesting that microglial absence leading to non-glial scar related regulation during regeneration. Similar to the brain, zebrafish and medaka heart regeneration is also impaired and failed to resolve the inflammation in the absence of macrophages (Lai et al., 2017), equivalent of microglia. In rodents, microglia promote neurogenesis in the early postnatal forebrain SVZ (Shigemoto-Mogami et al., 2014) and are required for cardiac repair after ischemia (Aurora et al., 2014; Lorchner et al., 2015). In contrast, microglia are not required for axonal regeneration after spinal cord injury in zebrafish or for optic nerve regeneration in mouse (Hilla et al., 2017; Themistoklis M. Tsarouchas et al., 2018), where macrophages are required. Thus, while we find that activated microglia are critical for brain repair in zebrafish, their importance for regeneration appears to differ based upon CNS region, species or in non-neural tissues.

### Microglia are required for robust aIPC amplification and neurogenesis after injury

Our data indicate that microglial ablation reduces aIPC amplification and neurogenesis, which in turn impairs restoration of injured brain tissue. A live imaging study of neurogenesis in the intact and injured adult zebrafish brain demonstrated that zebrafish use symmetric cell divisions of Sox2 positive aIPCs to amplify the daughter cells and induce robust neurogenesis after injury (Barbosa et al., 2015). Such NPC symmetric divisions do not occur in the intact brain, suggesting that this specific type of cell division promotes injury-induced proliferation and neurogenesis, while the aNSC divides only once in both the intact and injured brain (Barbosa et al., 2015). In our model, microglial ablation specifically suppressed aIPCs proliferation, indicating that microglia stimulate brain repair by specifically expanding this population for robust regeneration. In the mammalian brain, microglia accumulate near the stem cell niches in the SVZ and hippocampus during development to maintain NPC proliferation, and blocking the chemokine CX3CL2 or ablating microglia negatively affects progenitor cells in the cortex (Arnò et al., 2014). Microglial IGF1 and TGF-ß are also necessary to induce NPC proliferation after seizures or brain ischemia (Ali et al., 2015; Choi et al., 2017; Choi et al., 2008). In addition, microglia actively remodel adult hippocampal neurogenesis by maintaining the NPC population in mice (Diaz-Aparicio et al., 2020; Sierra et al., 2010). These reports are consistent with our findings that microglia regulate NPCs during reparative neurogenesis.

### Microglia-induced Tnfα activates Stat3 and ß-Catenin signaling and NPC proliferation

Stat3 and β-Catenin signaling are necessary for aNSC/NPC proliferation in many tissues. Our work indicates that microglial activation is necessary to stimulate these signaling factors to initiate the regenerative response. Stat3 is necessary for fin, retina and heart regeneration in fish and other animals (Y. Fang et al., 2013; Liang et al., 2012; Nelson et al., 2012; Sun et al., 2011; Zhao et al., 2014). For example, Stat3 activates retinal Mueller glia to promote regeneration, and Stat3 inhibition blocks both retina and heart repair (Nelson et al., 2012; Zhao et al., 2014). Jak/Stat and Wnt/ß-Catenin signaling promotes stem cell self-renewal (Goldman, 2015; Richmond et al., 2018; Shimizu et al., 2018; Todd et al., 2016; Zhao et al., 2014) and blocking negative regulators of Jak/Stat signaling or treating with Wnt/ß-Catenin inhibitors is sufficient to stimulate cell proliferation in uninjured tissue (Goldman, 2015; Hao et al., 2006; Ramachandran et al., 2011). Notably, Stat3 and ß-Catenin are induced by many inflammatory cytokines that are secreted by microglia during inflammation (Liu et al., 2013; Mori et al., 2011; Wan et al., 2014), consistent with our findings in brain regeneration.

We show that activated microglia are necessary for *tnfα* expression that stimulates Stat3 and ß-Catenin signaling to promote regeneration (Fig. 8p). Only 20% of microglia expressed *tnfα* in our model. Nonetheless, microglial signaling is likely required for most, if not all, of the *tnfα* expression after injury, as their suppression markedly reduces *tnfα* levels. Our work also suggests that Tnfα is downstream of microglial signaling, as inhibition of Tnfα drastically reduces regeneration without changing the microglial population. Although considered to be a pro-inflammatory cytokine, Tnfα is necessary for inducing regeneration after damage to the zebrafish brain, retina, and spinal cord (HH, 2018; Nelson et al., 2013; M. Nguyen-Chi et al., 2017; Mai Nguyen-Chi et al., 2017; Roehl, 2018; Themistoklis M. Tsarouchas et al., 2018).

Repressing notch signaling and expressing *tnfα* is sufficient to induce Muller glial proliferation in retina (Conner et al., 2014). Microglial Tnfα also induces NPC proliferation after stroke (Iosif et al., 2006) and is necessary for hematopoietic regeneration in murine cells (Bowers et al., 2018). Consistent with our proposed interaction between Tnfα and stat3/ß-catenin signaling after brain injury (Fig. 8p), recent work suggests that Tnfα activates ß-Catenin signaling during hair follicle regeneration (Wang et al., 2017), and microglial-stimulated Il1b and Tnfα induce Stat3 signaling in inflammatory arthritis (Mori et al., 2011). Manipulating Tnfa specifically in microglia or in neural stem cells and temporally will provide more mechanistic details during regeneration.

### Macrophages and neutrophils are not sufficient for brain regeneration

Our study shows that presence of macrophages and neutrophils are not necessary or sufficient for successful regeneration. In contrast, neutrophils might be inhibiting the regeneration process by extending inflammation longer than the control. In the absence of microglia, macrophages are present in the clodronate brain and neutrophils are accumulated more than the control, both in clodronate brain and the genetic mutants. In contrast, Macrophages are necessary for spinal cord repair, while 4C4 microglia are dispensable(T. M. Tsarouchas et al., 2018) and not necessary for axonal regeneration(Hilla et al., 2017). We speculate this difference is specific to neural tissues and CNS, compared to peripheral nervous system and other tissues, where macrophages are necessary for regeneration(Lai et al., 2017). However, both in the brain and spinal cord, neutrophils accumulate in *irf8* and *csf1rdm* mutants, and inhibits regeneration via manipulating inflammation resolution like in the brain (T. M. Tsarouchas et al., 2018).

### Activated microglia are critical for inflammation resolution

We found that activated microglia accumulate closer to the injury zone than infiltrating neutrophils at 2 dpl, the peak of microglial activation after injury. Microglial ablation leads to persistent neutrophil accumulation at the lesion, suggesting that activated microglia regulate neutrophil persistence/clearance after brain injury. *tnfα* is strongly expressed for the first 24 hpl and then decreases at 48 hpl, both in our model and during fin regeneration (Mai Nguyen-Chi et al., 2017). In the absence of microglia, we observed persistently elevated *tnfα* expression, likely arising from the accumulation of *tnfα*-expressing neutrophils, suggesting that the neutrophils extend the proinflammatory state. Similarly, microglial ablation after zebrafish cardiac injury led to the accumulation of neutrophils and impaired regeneration (de Preux Charles et al., 2016; Lai et al., 2017). In Medaka, cardiac regeneration was impaired with delayed macrophage recruitment and neutrophil accumulation, but inducing early macrophage recruitment improved regeneration, neutrophil clearance, and scar resolution (Lai et al., 2017). Furthermore, inducing neutrophil apoptosis resolved inflammation in zebrafish during tailfin regeneration (Hoodless et al., 2016), and in mice, microglia prevented the accumulation of neutrophils to suppress inflammation in an Alzheimer’s disease model (Unger et al., 2018). All these results are in line with our finding that microglial recruitment is crucial for removing pro-inflammatory neutrophils to resolve injury-induced inflammation.

Taken together, our data show that microglia are necessary for successful brain regeneration, and that microglial stimulation of Tnfα activates Stat3 and ß-Catenin signaling to promote aIPC amplification and neurogenesis. Also, macrophages are not sufficient for regeneration. Notably, our findings indicate that activated microglia attenuate inflammation, in part by clearing the infiltrating neutrophil population. Hence, the current study underscores the need to delineate specific inflammatory processes that promote regeneration in zebrafish, as well as those that inhibit regeneration in the mammalian brain. Further study to identify the microglial signaling molecules responsible for resolving inflammation in zebrafish should provide candidates to test in rodent injury models and lead to strategies for repairing the injured mammalian brain.

### Experimental Procedures

#### Animal care and transgenic fish lines

Fish were kept under standard conditions in accordance with guidelines and approval of the Institutional Animal Care & Use Committee at the University of Michigan as previously described (Skaggs et al. 2014). AB lines were used as wild type and the following transgenic lines or mutants were used in this study: *Tg*(*gfap*:GFP) (Ramachandran et al., 2011) and *Tg*(*olig2*:GFP) (provided by Bruce Appel, University of Colorado Denver, (Park et al., 2007)), *Tg* BAC(*tnfa*:GFP) (Marjoram et al., 2015), Tg(*1016tubulin1a*:GFP) (Fausett & Goldman, 2006), *Tg*(*gfap*:stat3-GFP) (Zhao & Goldman, 2014), *Tg* (*ascl1a*:GFP) (Wan et al., 2012), *Tg*(tcf7miniP:dGFP) (Shimizu et al., 2012), *Tg*(*mpeg1*:GFP) (F. Ellett et al., 2011) gl22, *Tg*(*mpeg1*:mcherry)(Felix Ellett et al., 2011) gl23, *Tg*(*hsp70*l:*ascl1a*) (Elsaeidi et al., 2018), *Tg*(*her4.1*:CreER^T2^) (Boniface et al., 2016), *Tg*(*ß-actin2*:loxP-mCherry-loxP-GFP) (Ramachandran et al., 2010), *Tg*(*mfap4*:dLanYFP-CAAX)(Walton et al., 2015), *Tg*(*mfap4*:TdTomato)(Walton et al., 2015), *irf8^st95 (Shiau et al., 2015)^*, *csf1r^dm^* (Oosterhof et al., 2018).

#### Generating transgenic zebrafish

ß-Catenin activity is regulated by protein stability. The stability is controlled by phosphorylation at Ser33, Ser37, Thr41 and Ser45, by tagging them for degradation (V. S. Li et al., 2012; MacDonald et al., 2009). We created the constitutively active protein by mutating these sites (S33A, S37A, T41A and S45A) by site-directed mutagenesis to block their degradation (Baba et al., 2006; Barth et al., 1997). In Contrast, Stat3 phosphorylation at Y705 stimulates its activation by dimerization (Bromberg et al., 1999; Levy & Darnell, 2002). By forcing dimerization using cysteine substitutions in its carboxyl terminus (A662C and N664C), we created a constitutively active Stat3 C terminus that does not require Y705 phosphorylation. *Tg* (*hsp70l*:CA*-ß-catenin*) and *Tg* (*hsp70l:*CA-*stat3*) lines were created using standard recombinant DNA technology using Tol2 vector backbone. Expression constructs were injected into the embryo as previously reported (Fausett & Goldman, 2006)

#### Brain injury model, mitotic labeling, 4-OH-TMX treatment, RNA isolation and real-time PCR

Brain injury, EdU intra-peritoneal injection, EdU pulse-chase, 4-OH-TMX treatment, were performed as previously described (Skaggs et al., 2014). Male and female fish (6-8 months old) were used for the experiments in this study, unless reported otherwise. RNA was isolated from the telencephalon using RNeasy kit (Qiagen). Total RNA was used for cDNA synthesis using iScript kit and real-time qRT-PCR (quantitative real-time PCR) were done in triplicate with AB PowerSYBR green mix on an iCycler (Bio-Rad). The primers used for qRT-PCR are listed in Table.

#### Pharmacological experiments

Fish were treated with following compounds: 1 mM Tnfa protein synthesis inhibitor (Thalidomide), and 500 nM PLX3397, a Csf1r inhibitor. Thalidomide was injected directly into the brain during injury and PLX3397 was delivered through fish water beginning 7 days before injury through 2 days after injury. The fish water was replaced daily during PLX3397 treatment, and the fish were fed every day for two hours. DMSO (0.05%) was used as the vehicle control for PLX3397 experiments.

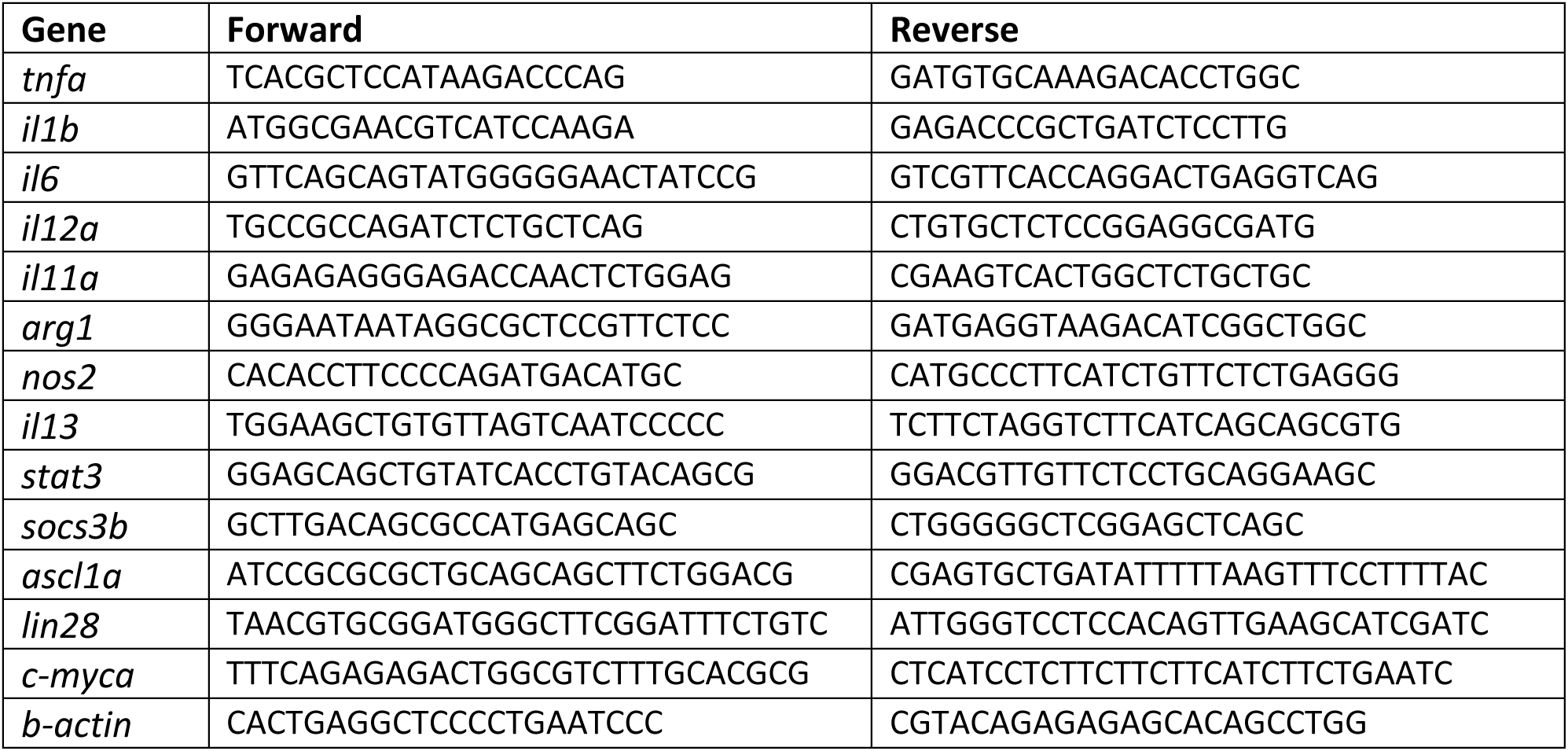

#### Clodronate liposome treatment

A 2.5 μl solution containing 5 mg/ml liposomes carrying clodronate in PBS or PBS only (Liposoma Research) was injected into the right telencephalon during brain lesioning using a Hamilton syringe with 30G needle. For tracking liposomes, we used a fluoro-liposome kit (Encapsula) in which liposomes carry a red fluorescent marker. The fluoro-liposome solution was delivered as described above and fluorescence was imaged in brain sections after fixation.

#### Heat-shock treatment

Heat shock was done by immersing the fish in pre-heated water at 37 C for 2 h followed by return to their regular temperature. The heat shock was done for 2 h periods three times per day after injury.

#### Tissue preparation and histological assays

Tissue preparation, immunohistochemistry, EdU (5-ethynyl-2’-deoxyuridine) labeling and TUNEL (Terminal deoxynucleotidyl transferase dUTP nick end labeling) were performed as described previously (Skaggs et al., 2014). Primary antibodies used were mouse anti (α)-4C4 (1:250, provided by Peter Hitchcock, University of Michigan, Ann Arbor), mouse α-glutamine synthetase (1:500, Chemicon MAB302), mouse α-HuC/D (1:250, Molecular Probes A-21271), rabbit α-GFP (1:2000, Molecular Probes A-11122), mouse α-chondroitin sulfate (1:50, Abcam AB11570), rabbit α-Gfap (1:250, Dako Z0334), rabbit α-Mpx (1:200), mouse α-mcherry (1:500), mouse α-PCNA (1:500 with antigen retrieval, Dako M087901), rabbit α-L-Plastin (1:1000, provided by Ji Feng, University of Edinburgh, UK), rabbit α-Sox2 (1:500, Abcam AB97959). Secondary antibodies used were α-mouse or -rabbit Alexa Fluor 488-conjugate raised in goat (1:300, Invitrogen A-11001 or A-11008, respectively), and α-mouse or rabbit Alexa Fluor 594-conjugate raised in goat (1:300, Invitrogen A-11032 or A-11037, respectively). Bisbenzimide was used for nuclear staining.

#### Microscopy and quantification of immunohistochemistry

Images were obtained using a Leica DMI 6000B epifluorescence microscope equipped with a Hamamatsu digital camera or on an Olympus FLUOVIEW FV1000 inverted confocal microscope. Images were processed using ImageJ Fiji plugin. For stacked confocal images, light attenuation was adjusted using the Stack Contrast Adjustment plugin. For high-resolution confocal images, stitching was done using the Stitch Grid of Images plugin. All quantifications were done using a minimum of five brains per condition and averaging across at least five 12-µm sections per brain located approximately 60 µm apart. All images were blinded to the observer for quantification. Any brightness or contrast changes were applied uniformly across all images for a given quantification. All normalizations to telencephalic area or VZ length were measured in ImageJ.

For calculating GFP intensity, images were quantified using Fiji software (Schindelin et al., 2012). Images were used to measure Raw Integrated Density of the selected region of interest after normalization of the background reading using three selected regions with no visible staining. The corrected fluorescence intensity was calculated using the following formula: Corrected total cellular fluorescence (CTCF) = Integrated density – (Area of selected cell x Mean fluorescence of background readings). The whole right telencephalon or left telencephalon were selected to quantify the intensity and multiple images at different depths of the section were included from a minimum of 5 different animals for quantification. The CTCF from the clodronate injured brain was compared to the control.

For microglial immunoreactivity, the average background grey value was multiplied by 1.5 and set as a threshold. Area past threshold stained for m4C4 was then normalized to total telencephalic area. The area of microglial immunoreactivity, EdU-positive cell numbers, TUNEL-positive cell numbers and counts of other immunoreactive cell numbers were all calculated using ImageJ software. For persistent tissue disruption at 90 dpl, bisbenzimide-stained tissues were blindly scored for the presence of altered tissue morphology, such as a cavity or other tissue disruption consistent with a persistent lesion. To be conservative, only brains with obvious disruptions were counted, and all others were considered as fully regenerated.

#### Statistical analyses

All statistical analyses were conducted using GraphPad or Prism 8.0. ANOVA, Chi-square and t-test were used to analyze statistical difference between groups. All the experiments were done in triplicates and 5 animals per experiment was used, unless stated differently. All values are displayed as mean ± standard error of mean, unless otherwise stated in the figure legends. In graphs, p-values are represented as follows: ns - not significant (p-value > 0.05); *, p-value ≤ 0.01; **, p-value ≤ 0.001; ***, p-value ≤ 0.0001.

## Acknowledgements

This work is supported by the University of Michigan, the Global ischemia foundation (K.P. and J.P.) and a National Science Foundation graduate research fellowship (J.C.). We would like to thank Dr. Michel Bagnat for providing the BAC (*tnfa*:GFP) fish, Dr. Bruce Appel for providing the Tg(*olig2*:GFP) fish, Dr. Peter Hitchcock for providing 4C4 antibody, Dr. David Tobin for providing *mafp4* reporter lines, Dr. Celia Shiau for providing *irf8* mutant fish, Dr. Tjakka van Ham for providing *csf1r*^dm^ fish, Dr. Diana M. Mitchell for providing mpeg1:mcherry fish, and Dr. Yi Feng for providing the L-Plastin antibody. We also would like to thank Muchu Zhou and Zachary Rakowski for assistance with fish care and maintenance.

## Author contributions

J.P. D.G, and P.K designed the study. P.K, J.C, K.S, Y.Q, M.C, N.C, J.R, V.K, A.P, D.A, and C.P. conducted the experiments, collected data, and contributed to data analysis. P.K, and J.P wrote the manuscript.

## Competing interests

The authors declare no competing interests.

## Supplemental Figure Legends

**Figure S1.**
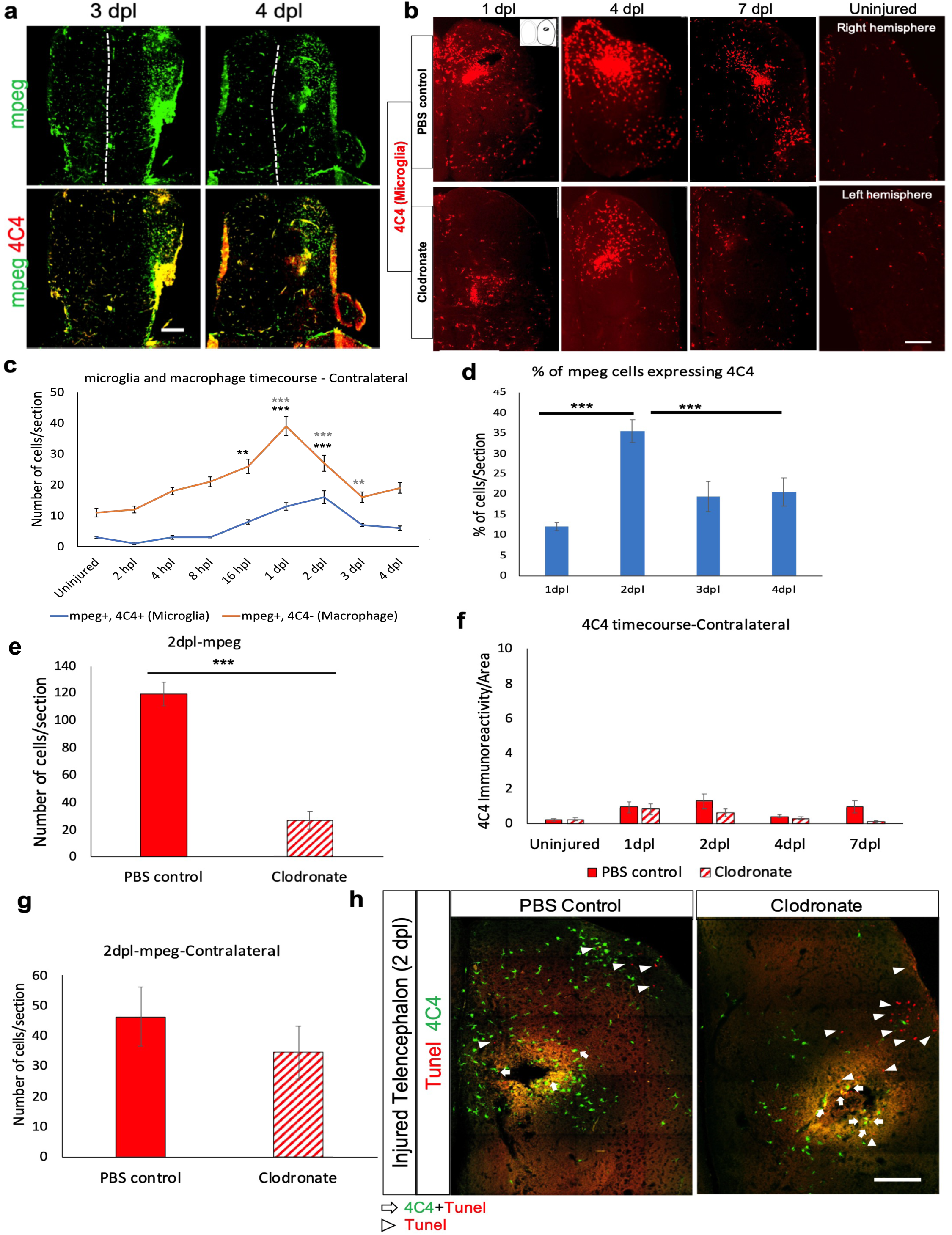
Liposomal clodronate efficiently suppresses activated microglia. a. Confocal images of telencephala after reporter labeling for macrophages *tg*(*mpeg*:GFP; green; top panels) alone or with co-immunolabeling for activated microglia (4C4; red; bottom panels) at 3 and 4 dpl. The dashed line indicates the midline. b. 4C4 immunostaining of ipsilesional hemi-telencephala from control (top row) and clodronate-treated fish (bottom row) at 1, 4 or 7 dpl. The far right panels display 4C4 immunolabeling of uninjured hemi-telencephala. c. Quantification of microglia represented by blue line (*mpeg:*GFP+, 4C4+ double-positive) or macrophage represented by redline (*mpeg:*GFP+, 4C4-) from the contralateral telencephala at different time points after injury and in uninjured fish The asterisks indicate statistical significance compared to uninjured. **p<0.001, ***p<0.0001. d. Quantification of the percentages of *mpeg*:GFP- labeled cells expressing 4C4 at different time points. ***p<0.0001. e. Quantification of *mpeg*:GFP+ cells in the ipsilesional hemisphere at 2 dpl in control and clodronate groups. ***, p < 0.0001. f. Quantification of 4C4 immunoreactivity from uninjured fish and the contralateral hemi-telencephala at different time points after injury. g. Quantification of *mpeg*:GFP+ cells from the contralateral hemisphere at 2 dpl in control and clodronate groups. h. Confocal images of the injured hemi-telencephala after double labeling for 4C4 (green) and TUNEL stain (red) at 2 dpl in control liposome or clodronate liposome treated fish. Arrows shows double- positive cells and arrowheads shows cells positive for TUNEL stain only. Scale bars = 100 μm and the error bars indicate SEM.

**Figure S2.**
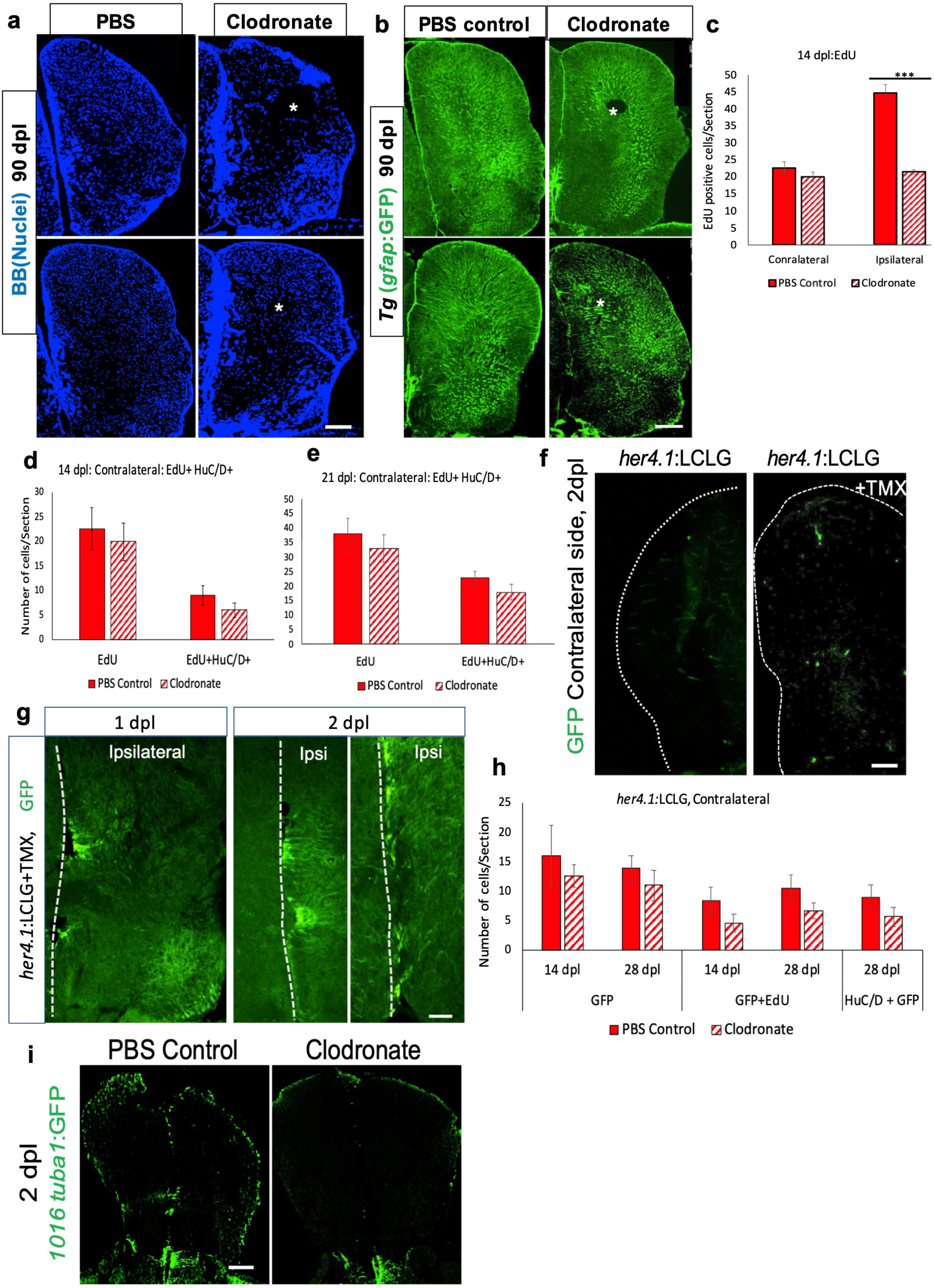
Microglial ablation impairs brain regeneration. a. Nuclear staining with bisbenzimide (BB) from two injured hemi-telencephala per group at 90 dpl after control (PBS) liposome or clodronate liposome treatment. Asterisks indicate the injury site. b. GFP immunolabeling in two ipsilesional hemi-telencephala per group from *gfap*:GFP reporter fish shows the glial response at 90 dpl in control (left panels) and clodronate (right panels) groups. Asterisks indicate the injury site. c. Quantification of EdU+ cells from both ipsilateral and contralateral telencephala at 14 dpl in control and clodronate groups. ***, p <0.0001. d, e. Quantification of EdU+ cells and EdU/HuC/D double-labeled cells from the contralesional side at 14 (d) and 21 (e) dpl. f. GFP immunolabeling of the contralesional hemispheres from *her4.1*:GFP reporter fish at 2 dpl with (right) or without (left) 4-OH-TMX treatment. The dotted lines outline the hemispheres. g. GFP immunolabeling of telencephala from *her4.1*:GFP reporter fish at 1 and 2 dpl after 4-OH-TMX treatment. Dotted lines denote the midline, and the injured hemisphere is on the right. h. Quantification of cells labeled for GFP, GFP+EdU and GFP+HuC/D in the contralateral hemi-telencephala at 14 and 28 dpl in control liposome or clodronate liposome treated fish. i. GFP immunostaining shows reporter activity from the injury responsive promoter tg(*1016tubulin1*:GFP) at 2 dpl in control and clodronate groups. Scale bars = 100 μm and error bars indicate SEM.

**Figure S3.**
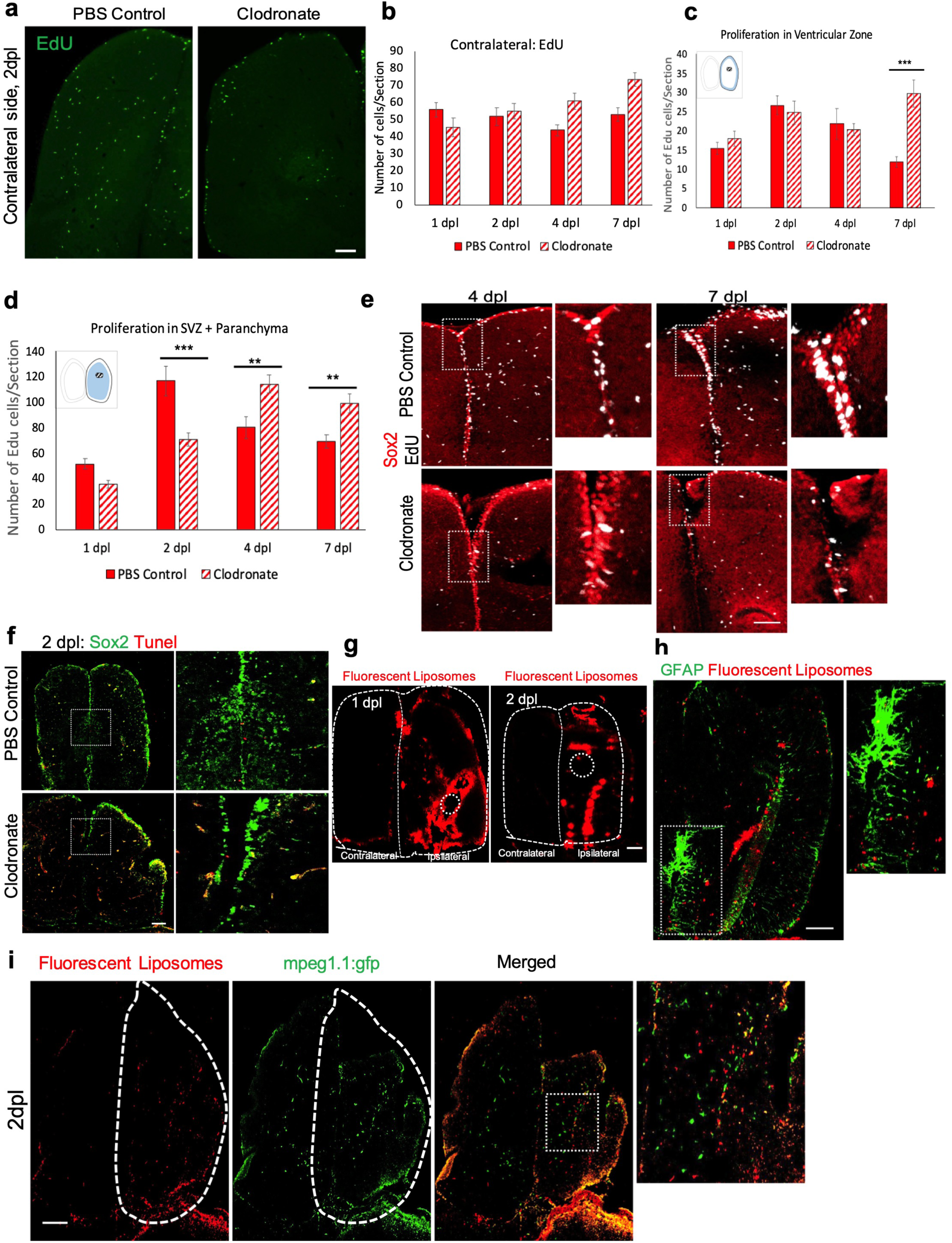
Microglial ablation inhibits progenitor cell amplification. a. EdU labeling of the contralateral side at 2 dpl in representative hemi-telencephala from control and clodronate groups. b. Quantification of EdU+ cells from the contralateral hemi-telencephala at 1, 2, 4 and 7 dpl. c. Quantification of EdU+ cells from the ventricular zone (blue outline in inset) of the ipsilesional hemi-telencephala at 1, 2, 4 and 7 dpl. ***, p<0.0001. d. Quantification of EdU+ cells in the subventricular zone (SVZ) and the parenchymal region (blue label in inset) of the ipsilesional hemi-telencephala at 1, 2, 4 and 7 dpl. **, p <0.01; ***, p<0.0001. e. Confocal images from the anterior midline telencephala show double labeling for Sox2 and EdU in control (top row) and clodronate groups (bottom row) at 4 and 7 dpl. EdU was administered at 2 dpl. Boxed areas are shown at higher magnification on the right of each pair of images. f. Confocal images displaying Sox2 and TUNEL double labeling at 2 dpl in control and clodronate groups. The right-sided panels show higher magnification views of the boxed regions. g. Images show red fluorescent liposomes at 1 and 2 dpl. The telencephala are outlined by dotted lines and the circles mark the injury sites. h. Double labeling for radial glia with GFAP immunolabeling (green) and red fluorescent liposomes at 2 dpl. The boxed area is shown at higher magnification on the right. Note minimal uptake of red fluorescent liposomes by radial glia. i. Staining for macrophages (*mpeg*:GFP) and red fluorescent liposomes at 2 dpl. The dashed lines outline the injured hemi-telencephala and the boxed area in the merged image is shown at higher magnification on the right. Scale bars = 100 μm and the error bars indicate SEM.

**Figure S4.**
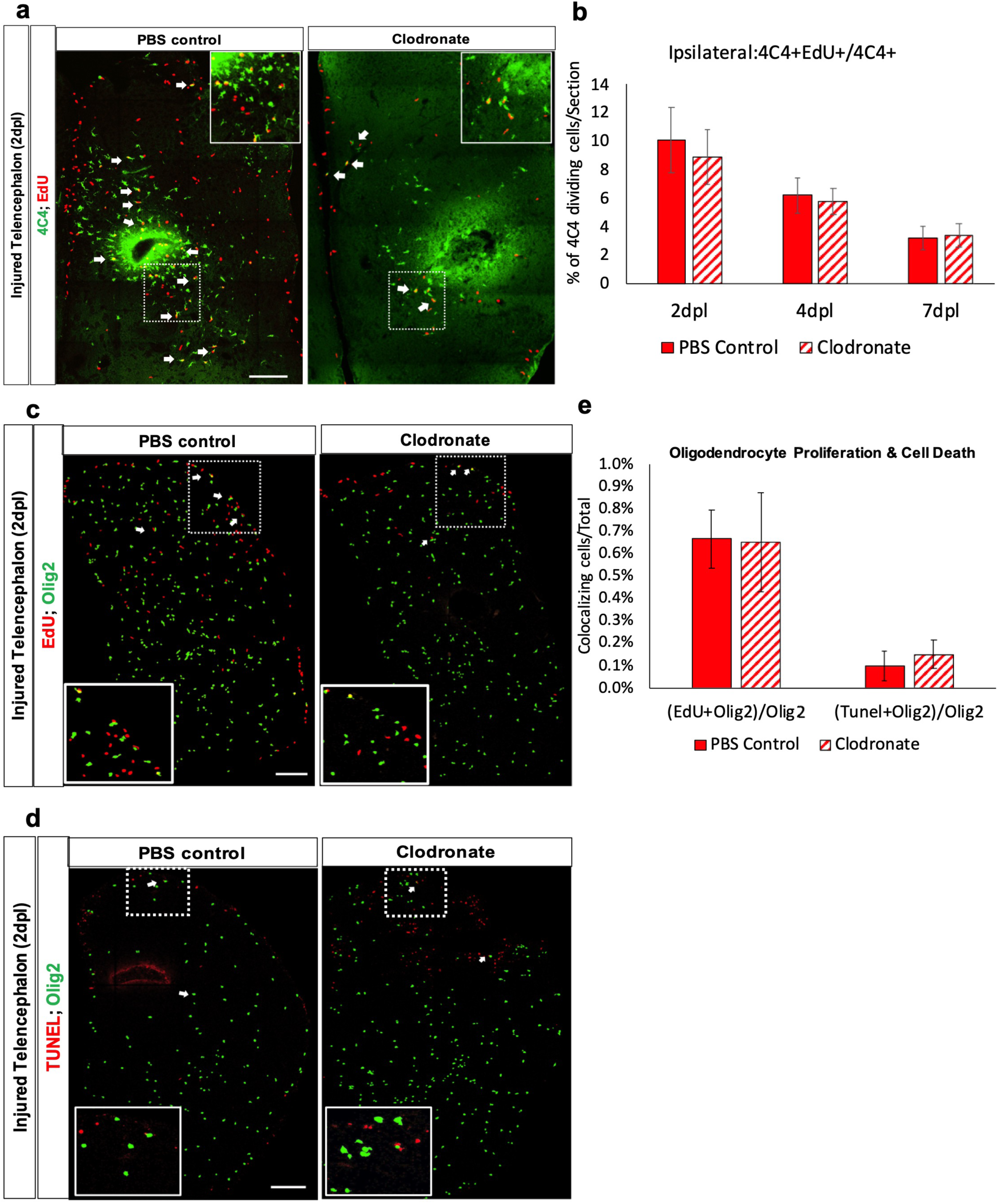
Microglial and oligodendrocyte proliferation after injury. a. Double labeling for 4C4 and EdU in the ipsilateral hemi-telencephala at 2 dpl in control and clodronate groups. Arrows indicate double-labeled cells. The insets at upper right show higher magnification views of the boxed areas. b. Quantification of the percentage of 4C4+ cells at 2, 4 and 7 dpl in the injured hemi-telencephala of control and clodronate groups that had incorporated EdU at 2 dpl. c. Double labeling for Olig2 and EdU in the ipsilateral hemi-telencephala at 2 dpl. The insets at bottom left show higher magnification views of the boxed areas. d. Double labeling for Olig2 and TUNEL at 2 dpl from the ipsilesional side in control and clodronate groups. Arrows indicate Olig2/TUNEL double-positive cells. The insets at bottom left show higher magnification views of the boxed areas. e. Quantification of the percentages of proliferating (EdU+) and apoptotic (TUNEL+) Olig2+ cells. Scale bars = 100 μm and error bars indicate SEM.

**Figure S5.**
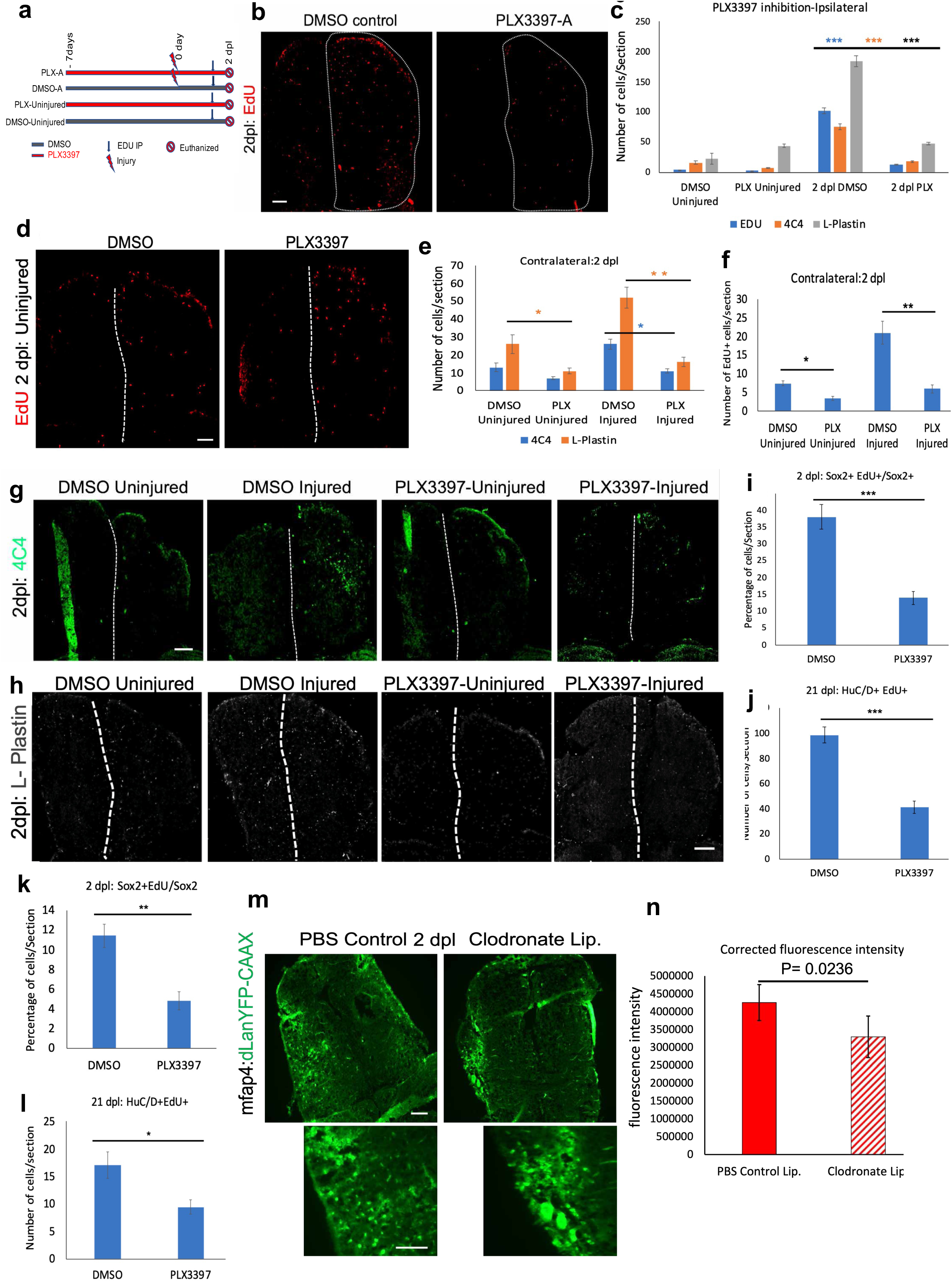
Csf1r inhibition with PLX3397 blocks injury-induced progenitor cell proliferation and adult neurogenesis. a. Schematic of the experimental design and time course. Red lines are PLX3397 (PLX)-treated groups and blue lines represent DMSO-treated controls. b, g, h. Confocal images of immunolabeling to identify activated microglia (4C4; g), leukocytes (L-Plastin; h) and proliferating cells (EdU; b) at 2 dpl in telencephala from uninjured or injured fish treated with DMSO or PLX3397. Dashed lines indicate the midline in b, c and the injured hemi-telencephela are outlined in b. For d, EdU was administered at 2 dpl after a 9-day exposure to PLX3397 or DMSO and 2 hours prior to euthanasia. c. Quantification of L-Plastin, 4C4 and EdU labeled cells at 2 dpl after treatment with DMSO or PLX3397. Note the increased numbers of labeled cells seen in the control lesioned group (2 dpl DMSO) but not in the PLX3397-treated, lesioned group (2 dpl PLX). ***, p<0.0001 by t-test. d. Confocal images of EdU labeling in the telencephala of uninjured fish treated with vehicle (DMSO) or PLX3397. The dotted lines denote the midline. e. Quantification of 4C4- and L-Plastin-positive cells in uninjured hemi-telencephala or at 2 dpl in fish treated with DMSO or PLX3397. *, p < 0.01; **, p <0.001. f. Quantification of EdU+ cells in the contralesional hemi-telencephala of fish treated vehicle (DMSO) or PLX3397 at 2 dpl or without injury. i, j. Quantification of EdU/Sox2 double-labeled cells at 2 dpl (f) and EdU/HuC/D double-labeled cells from the ipsilateral side at 21 dpl. k. Quantification of the percentage of Sox2+ cells that incorporated EdU in the contralesional side at 2 dpl. l. Quantification of HuC/D and EdU double-labeled cells in the contralesional side at 21 dpl. EdU was administered at 2 dpl. m. image of mfap4 reporter line for macrophages at 2 dpl after control or clodronate treatment. n. Quantification of mfap4 reporter intensity and corrected for area using imageJ. T- test was used to test the significance and error bar indicates SEM. Scale bar = 100 μm and error bars indicate SEM.

**Figure S6.**
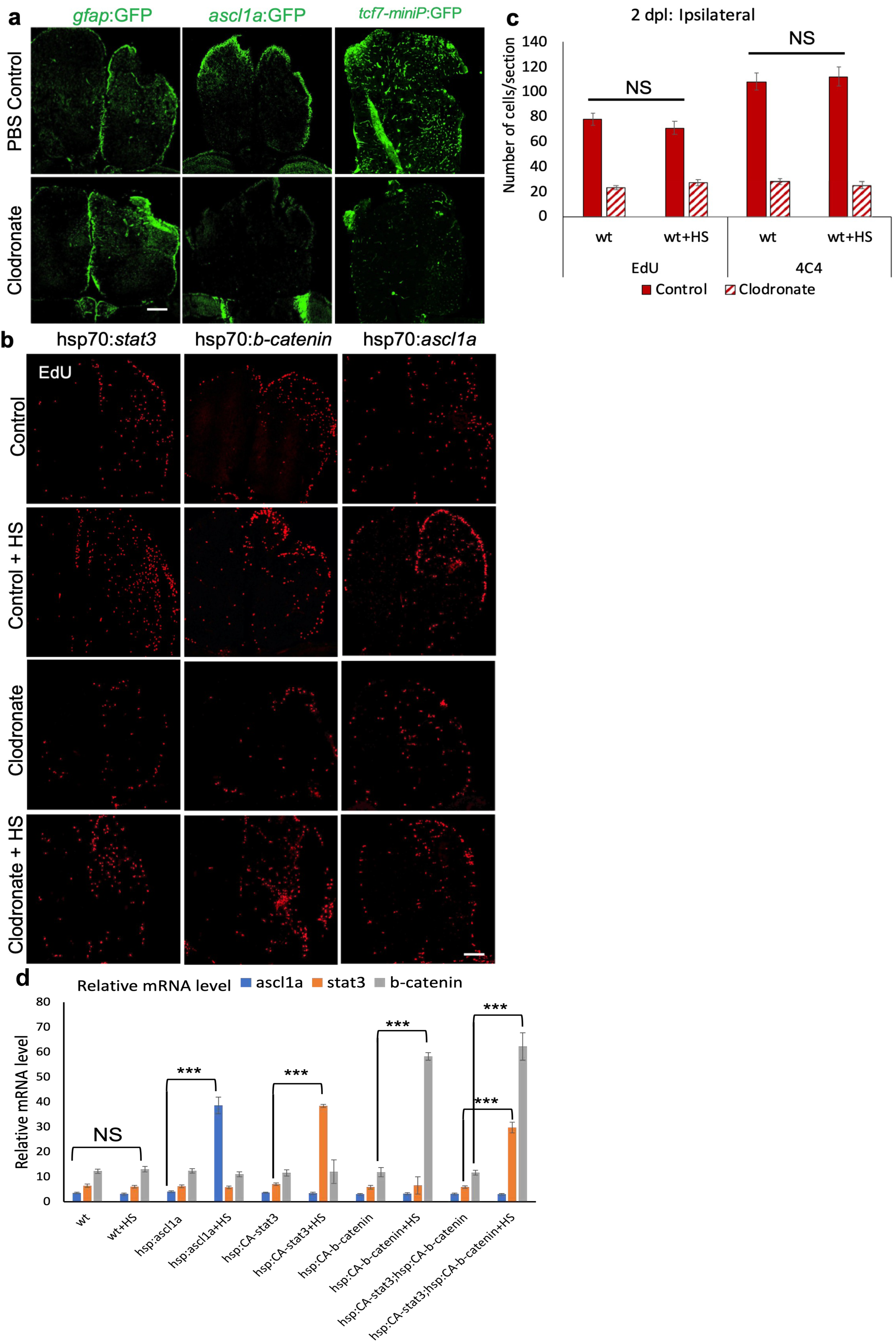
Microglia are necessary to induce pro-regenerative signaling after injury. a. GFP staining of reporter lines for *gfap*, *ascl1a* and Wnt signaling (tcf7) at 2 dpl in control and clodronate groups. b. EdU labeling at 2 dpl for rescue analysis with or without heat shock (HS) in both control liposome and clodronate liposome treated groups. HS activates the HS promoter (hsp) driving *stat3*, b-*catenin* or *ascl1a*. c. Quantification of EdU- or 4C4-labeled cells from wildtype brains at 2 dpl with or without HS shows that HS alone has no effect on cell numbers. d. qRT-PCR for ascl1a, stat3, b-catenin in the brain from overexpression by heat shock analysis. t-test and was used. ***, p = 0.0001. Scale bars = 100 μm and error bars indicate SEM except d.

**Figure S7.**
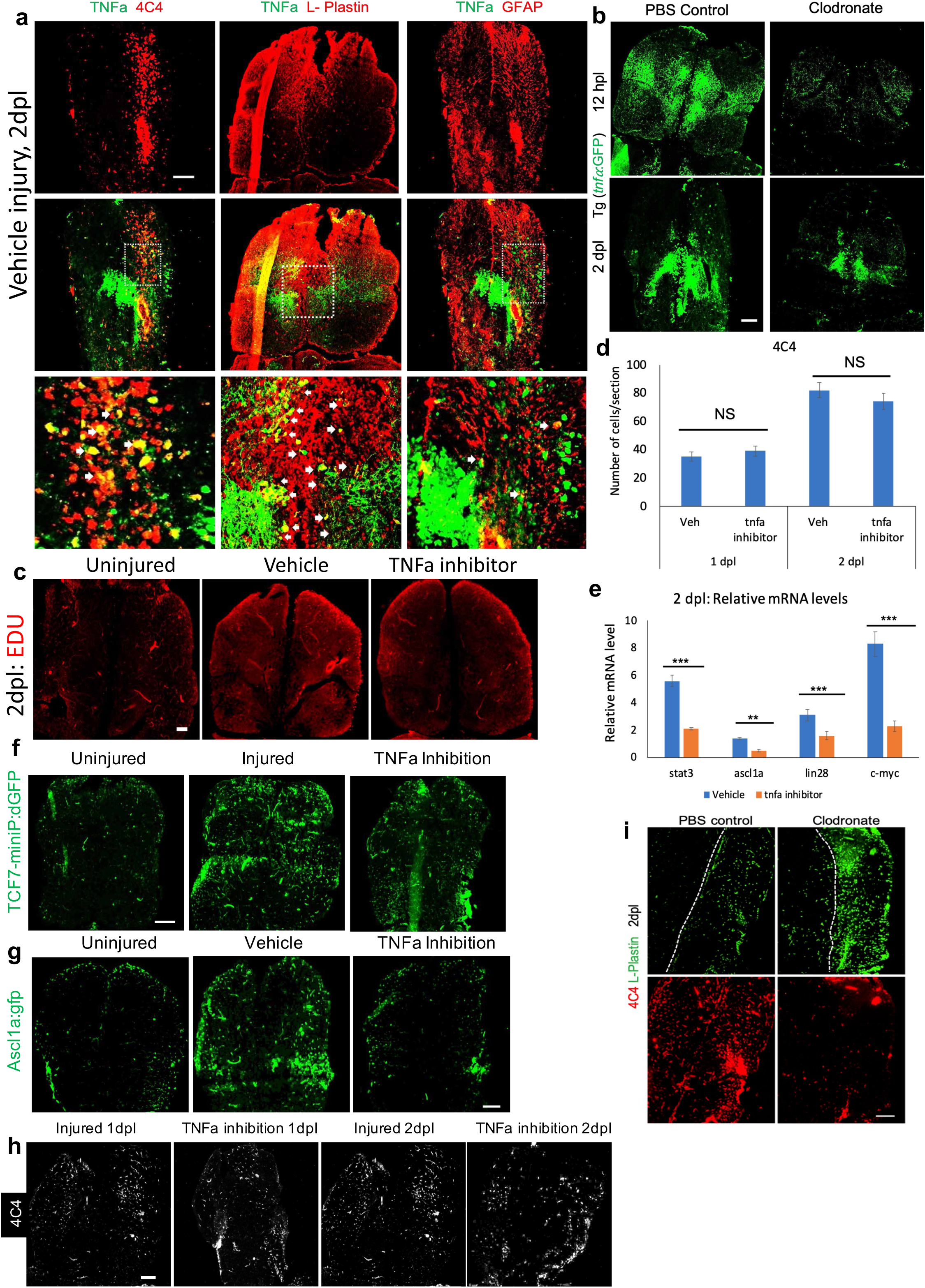
Microglial-dependent tnfa is necessary to activate regenerative signaling. a. Double labeling for tnfa (green) and 4C4, L-Plastin or gfap at 2 dpl after vehicle injury. The middle row shows merged images with the boxed regions displayed at higher magnification in the bottom row. b. Confocal images showing *tnfa*:GFP expression after control liposome or clodronate liposome lesioning at 12 hpl and 2 dpl. c. EdU labeling in the telencephala of an uninjured fish or fish at 2 dpl treated with vehicle or the tnfa inhibitor thalidomide. d. Quantification of 4C4+ cells in the injured hemi-telencephala of fish treated with vehicle (Veh) or thalidomide at 1 or 2 dpl. NS, not statistically significant. e. qRT-PCR showing relative mRNA levels of the indicated genes at 2 dpl from the ipsilateral hemi-telencephala of fish treated with vehicle or thalidomide. The error bars indicate SD. **, p = 0.001; ***, p = 0.0001. f. Wnt signaling-driven GFP reporter activity in the telencephala of an uninjured fish or at 2 dpl treated with vehicle or thalidomide. g. *ascl1a*-driven GFP reporter expression in the telencephala of an uninjured fish or fish at 2 dpl treated with vehicle or thalidomide. h. 4C4 immunoreactivity in the injured telencephala (right hemispheres) of fish at 1 or 2 dpl after treatment with vehicle or thalidomide. i. immunolabelling for 4C4 and L-Plastin at 2 dpl after control or clodronate treatment. Scale bars = 100 μm, and error bars indicate SEM.

## Notes

### Competing Interest Statement

The authors have declared no competing interest.

### Summary of Updates

The current version has two different genetic mutants to confirm our earlier findings and a few additional experiments to clarify the role of microglia, macrophage, and neutrophils. Also, we have added a few experiments to strengthen our claim from this paper. We have reorganized the figure to make it easy to read the story coherently.

## REFERENCES

Ali, I., Chugh, D., & Ekdahl, C. T. (2015). Role of fractalkine-CX3CR1 pathway in seizure-induced microglial activation, neurodegeneration, and neuroblast production in the adult rat brain. Neurobiol Dis, 74, 194–203. https://doi.org/10.1016/j.nbd.2014.11.009

Alunni, A., & Bally-Cuif, L. (2016). A comparative view of regenerative neurogenesis in vertebrates. Development, 143(5), 741–753. https://doi.org/10.1242/dev.122796

Alvarez-Buylla, A., & Garcia-Verdugo, J. M. (2002). Neurogenesis in adult subventricular zone. J Neurosci, 22(3), 629–634.

Alvarez-Buylla, A., & Garcıá-Verdugo, J. M. (2002). Neurogenesis in Adult Subventricular Zone.

Arnò, B., Grassivaro, F., Rossi, C., Bergamaschi, A., Castiglioni, V., Furlan, R., . . . Muzio, L. (2014). Neural progenitor cells orchestrate microglia migration and positioning into the developing cortex [Research]. Nature Communications, 5, 5611. https://doi.org/doi:10.1038/ncomms6611

Aurora, A. B., Porrello, E. R., Tan, W., Mahmoud, A. I., Hill, J. A., Bassel-Duby, R., . . . Olson, E. N. (2014). Macrophages are required for neonatal heart regeneration. J Clin Invest, 124(3), 1382–1392. https://doi.org/10.1172/jci72181

Baba, Y., T, Y., H, S., KP, G., S, H., & PW, K. (2006). Constitutively active beta-catenin promotes expansion of multipotent hematopoietic progenitors in culture. J Immunol, 177(4). https://doi.org/10.4049/jimmunol.177.4.2294

Barbosa, J. S., & Ninkovic, J. (2016). Adult neural stem cell behavior underlying constitutive and restorative neurogenesis in zebrafish. Neurogenesis (Austin), 3(1), e1148101. https://doi.org/10.1080/23262133.2016.1148101

Barbosa, J. S., Sanchez-Gonzalez, R., Giaimo, R. D., Baumgart, E. V., Theis, F. J., Götz, M., & Ninkovic, J. (2015). Live imaging of adult neural stem cell behavior in the intact and injured zebrafish brain. Science. https://doi.org/10.1126/science.aaa2729

Barth, A., AL, P., Y, A., KE, M., & WJ, N. (1997). NH2-terminal deletion of beta-catenin results in stable colocalization of mutant beta-catenin with adenomatous polyposis coli protein and altered MDCK cell adhesion. The Journal of cell biology, 136(3). https://doi.org/10.1083/jcb.136.3.693

Baumgart, E. V., Barbosa, J. S., Bally-Cuif, L., Gotz, M., & Ninkovic, J. (2012). Stab wound injury of the zebrafish telencephalon: a model for comparative analysis of reactive gliosis. Glia, 60(3), 343–357. https://doi.org/10.1002/glia.22269

Becker, C. G., & Becker, T. (2008). Adult zebrafish as a model for successful central nervous system regeneration. Restor Neurol Neurosci, 26(2-3), 71–80.

Becker, C. G., & Becker, T. (2015). Neuronal regeneration from ependymo-radial glial cells: cook, little pot, cook! Dev Cell, 32(4), 516–527. https://doi.org/10.1016/j.devcel.2015.01.001

Boniface, E. J., Lu, J., Victoroff, T., Zhu, M., & Chen, W. (2016). FlEx-based transgenic reporter lines for visualization of Cre and Flp activity in live zebrafish. https://doi.org/doi:10.1002/dvg.20526

Bosak, V., Murata, K., Bludau, O., & Brand, M. (2018). Role of the immune response in initiating central nervous system regeneration in vertebrates: learning from the fish. Int J Dev Biol, 62(6-7-8), 403-417. https://doi.org/10.1387/ijdb.180033vb

Bowers, E., Slaughter, A., Frenette, P. S., Kuick, R., Pello, O. M., & Lucas, D. (2018). Granulocyte-derived TNFα promotes vascular and hematopoietic regeneration in the bone marrow. Nat Med, 24(1), 95–102. https://doi.org/10.1038/nm.4448

Bromberg, J., MH, W., G, D., Y, Z., RG, P., C, A., & JE, D. (1999). Stat3 as an oncogene. Cell, 98(3). https://doi.org/10.1016/s0092-8674(00)81959-5

Chen, J., Poskanzer, K. E., Freeman, M. R., & Monk, K. R. (2020). Live-imaging of astrocyte morphogenesis and function in zebrafish neural circuits [OriginalPaper]. Nature Neuroscience, 1-10. https://doi.org/doi:10.1038/s41593-020-0703-x

Cheung, T. H., & Rando, T. A. (2013). Molecular regulation of stem cell quiescence. Nat Rev Mol Cell Biol, 14(6). https://doi.org/10.1038/nrm3591

Choi, J. Y., Kim, J. Y., Park, J., Lee, W. T., & Lee, J. E. (2017). M2 Phenotype Microglia-derived Cytokine Stimulates Proliferation and Neuronal Differentiation of Endogenous Stem Cells in Ischemic Brain. Exp Neurobiol, 26(1), 33–41. https://doi.org/10.5607/en.2017.26.1.33

Choi, Y. S., Cho, H. Y., Hoyt, K. R., Naegele, J. R., & Obrietan, K. (2008). IGF-1 Receptor-Mediated ERK/MAPK Signaling Couples Status Epilepticus to Progenitor Cell Proliferation in the Subgranular Layer of the Dentate Gyrus. Glia, 56(7), 791–800. https://doi.org/10.1002/glia.20653

Conner, C., Ackerman, K. M., Lahne, M., Hobgood, J. S., & Hyde, D. R. (2014). Repressing notch signaling and expressing TNFalpha are sufficient to mimic retinal regeneration by inducing Muller glial proliferation to generate committed progenitor cells. J Neurosci, 34(43), 14403–14419. https://doi.org/10.1523/jneurosci.0498-14.2014

Crain, J. M., Nikodemova, M., & Watters, J. J. (2013). Microglia express distinct M1 and M2 phenotypic markers in the postnatal and adult central nervous system in male and female mice. J Neurosci Res, 91(9), 1143–1151. https://doi.org/10.1002/jnr.23242

de Preux Charles, A. S., Bise, T., Baier, F., Marro, J., & Jazwinska, A. (2016). Distinct effects of inflammation on preconditioning and regeneration of the adult zebrafish heart. Open Biol, 6(7). https://doi.org/10.1098/rsob.160102

Diaz-Aparicio, I., Paris, I., Sierra-Torre, V., Plaza-Zabala, A., Rodriguez-Iglesias, N., Marquez-Ropero, M., . . . Sierra, A. (2020). Microglia Actively Remodel Adult Hippocampal Neurogenesis through the Phagocytosis Secretome. J Neurosci, 40(7), 1453–1482. https://doi.org/10.1523/JNEUROSCI.0993-19.2019

Ekdahl, C. T., Claasen, J. H., Bonde, S., Kokaia, Z., & Lindvall, O. (2003). Inflammation is detrimental for neurogenesis in adult brain. Proc Natl Acad Sci U S A, 100(23), 13632–13637. https://doi.org/10.1073/pnas.2234031100

Ekdahl, C. T., Kokaia, Z., & Lindvall, O. (2009). Brain inflammation and adult neurogenesis: the dual role of microglia. Neuroscience, 158(3), 1021–1029. https://doi.org/10.1016/j.neuroscience.2008.06.052

Ellett, F., Pase, L., Hayman, J. W., Andrianopoulos, A., & Lieschke, G. J. (2011). mpeg1 promoter transgenes direct macrophage-lineage expression in zebrafish. Blood, 117(4), e49–56. https://doi.org/10.1182/blood-2010-10-314120

Ellett, F., Pase, L., Hayman, J. W., Andrianopoulos, A., & Lieschke, G. J. (2011). mpeg1 promoter transgenes direct macrophage-lineage expression in zebrafish. https://doi.org/10.1182/blood-2010-10-314120

Elsaeidi, F., Macpherson, P., Mills, E. A., Jui, J., Flannery, J. G., & Goldman, D. (2018). Notch suppression collaborates with Ascl1 and Lin28 to unleash a regenerative response in fish retina, but not in mice. J Neurosci. https://doi.org/10.1523/jneurosci.2126-17.2018

Fang, Y., Gupta, V., Karra, R., Holdway, J. E., Kikuchi, K., & Poss, K. D. (2013). Translational profiling of cardiomyocytes identifies an early Jak1/Stat3 injury response required for zebrafish heart regeneration. https://doi.org/10.1073/pnas.1309810110

Fang, Y., Gupta, V., Karra, R., Holdway, J. E., Kikuchi, K., & Poss, K. D. (2013). Translational profiling of cardiomyocytes identifies an early Jak1/Stat3 injury response required for zebrafish heart regeneration. Proc Natl Acad Sci U S A, 110(33), 13416–13421. https://doi.org/10.1073/pnas.1309810110

Fausett, B. V., & Goldman, D. (2006). A role for alpha1 tubulin-expressing Müller glia in regeneration of the injured zebrafish retina. J Neurosci, 26(23), 6303–6313. https://doi.org/10.1523/JNEUROSCI.0332-06.2006

Fitch, M. T., & Silver, J. (2008). CNS injury, glial scars, and inflammation: Inhibitory extracellular matrices and regeneration failure. Exp Neurol, 209(2), 294–301. https://doi.org/10.1016/j.expneurol.2007.05.014

Gage, F. H. (2002). Neurogenesis in the Adult Brain. Journal of neuroscience. https://doi.org/10.1523/JNEUROSCI.22-03-00612.2002

Gerlach, J., Donkels, C., Munzner, G., & Haas, C. A. (2016). Persistent Gliosis Interferes with Neurogenesis in Organotypic Hippocampal Slice Cultures. Front Cell Neurosci, 10, 131. https://doi.org/10.3389/fncel.2016.00131

Ghosh, S., & Hui, S. P. (2016). Regeneration of Zebrafish CNS: Adult Neurogenesis. Neural Plast, 2016, 5815439. https://doi.org/10.1155/2016/5815439

Goldman, D. (2014). Muller glial cell reprogramming and retina regeneration. Nat Rev Neurosci, 15(7), 431–442. https://doi.org/10.1038/nrn3723

Goldman, D. (2015). Retinal Injury, Growth Factors, and Cytokines Converge on Î²-Catenin and pStat3 Signaling to Stimulate Retina Regeneration: Cell Reports. http://www.cell.com/cell-reports/abstract/S2211-1247(14)00724-4

Goldman, D., & Ding, J. (2000). Different regulatory elements are necessary for alpha1 tubulin induction during CNS development and regeneration. Neuroreport, 11(17), 3859–3863.

Goldman, D., Hankin, M., Li, Z., Dai, X., & Ding, J. (2001). Transgenic zebrafish for studying nervous system development and regeneration. Transgenic Res, 10(1), 21–33.

Götz, M., & Huttner, W. B. (2005). The cell biology of neurogenesis. Nat Rev Mol Cell Biol, 6(10), 777–788. https://doi.org/10.1038/nrm1739

Hao, J., Li, T. G., Qi, X., Zhao, D. F., & Zhao, G. Q. (2006). WNT/beta-catenin pathway up-regulates Stat3 and converges on LIF to prevent differentiation of mouse embryonic stem cells. Dev Biol, 290(1), 81–91. https://doi.org/10.1016/j.ydbio.2005.11.011

HH, R. (2018). Linking wound response and inflammation to regeneration in the zebrafish larval fin. Int J Dev Biol., 62(6-7-8), 473-477. https://doi.org/10.1387/ijdb.170331hr

Hilla, A. M., Diekmann, H., & Fischer, D. (2017). Microglia Are Irrelevant for Neuronal Degeneration and Axon Regeneration after Acute Injury. J Neurosci, 37(25), 6113–6124. https://doi.org/10.1523/jneurosci.0584-17.2017

Hoodless, L. J., Lucas, C. D., Duffin, R., Denvir, M. A., Haslett, C., Tucker, C. S., & Rossi, A. G. (2016). Genetic and pharmacological inhibition of CDK9 drives neutrophil apoptosis to resolve inflammation in zebrafish in vivo. Sci Rep, 5, 36980. https://doi.org/10.1038/srep36980

Huang, T., Cui, J., Li, L., Hitchcock, P. F., & Li, Y. (2012). The role of microglia in the neurogenesis of zebrafish retina. Biochemical and Biophysical Research Communications, 421(2), 214–220. https://doi.org/https://doi.org/10.1016/j.bbrc.2012.03.139

Iosif, R. E., Ekdahl, C. T., Ahlenius, H., Pronk, C. J., Bonde, S., Kokaia, Z., . . . Lindvall, O. (2006). Tumor necrosis factor receptor 1 is a negative regulator of progenitor proliferation in adult hippocampal neurogenesis. J Neurosci, 26(38), 9703–9712. https://doi.org/10.1523/jneurosci.2723-06.2006

Jassam, Y. N., Izzy, S., Whalen, M., McGavern, D. B., & El Khoury, J. (2017). Neuroimmunology of Traumatic Brain Injury: Time for a Paradigm Shift. Neuron, 95(6), 1246–1265. https://doi.org/10.1016/j.neuron.2017.07.010

Jin, X., & Yamashita, T. (2016). Microglia in central nervous system repair after injury. J Biochem, 159(5), 491–496. https://doi.org/10.1093/jb/mvw009

Katz, S., Cussigh, D., Urban, N., Blomfield, I., Guillemot, F., Bally-Cuif, L., & Coolen, M. (2016). A Nuclear Role for miR-9 and Argonaute Proteins in Balancing Quiescent and Activated Neural Stem Cell States. Cell Rep, 17(5), 1383–1398. https://doi.org/10.1016/j.celrep.2016.09.088

Kizil, C., Kaslin, J., Kroehne, V., & Brand, M. (2012). Adult neurogenesis and brain regeneration in zebrafish. Dev Neurobiol, 72(3), 429–461. https://doi.org/10.1002/dneu.20918

Kizil, C., Kyritsis, N., Dudczig, S., Kroehne, V., Freudenreich, D., Kaslin, J., & Brand, M. (2012). Regenerative neurogenesis from neural progenitor cells requires injury-induced expression of Gata3. Dev Cell, 23(6), 1230–1237. https://doi.org/10.1016/j.devcel.2012.10.014

Kroehne, V., Freudenreich, D., Hans, S., Kaslin, J., & Brand, M. (2011). Regeneration of the adult zebrafish brain from neurogenic radial glia-type progenitors. Development, 138(22), 4831–4841. https://doi.org/10.1242/dev.072587

Kyritsis, N., Kizil, C., Zocher, S., Kroehne, V., Kaslin, J., Freudenreich, D., . . . Brand, M. (2012). Acute inflammation initiates the regenerative response in the adult zebrafish brain. Science, 338(6112), 1353–1356. https://doi.org/10.1126/science.1228773

Lai, S. L., Marín-Juez, R., Moura, P. L., Kuenne, C., Lai, J. K. H., Tsedeke, A. T., . . . Stainier, D. Y. (2017). Reciprocal analyses in zebrafish and medaka reveal that harnessing the immune response promotes cardiac regeneration. Elife, 6. https://doi.org/10.7554/eLife.25605

Levy, D., & Darnell, J. (2002). Stats: transcriptional control and biological impact. Nature reviews. Molecular cell biology, 3(9). https://doi.org/10.1038/nrm909

Li, L., Jin, H., Xu, J., Shi, Y., & Wen, Z. (2011). Irf8 regulates macrophage versus neutrophil fate during zebrafish primitive myelopoiesis. Blood, 117(4), 1359–1369. https://doi.org/10.1182/blood-2010-06-290700

Li, L., Yan, B., Shi, Y.-Q., Zhang, W.-Q., & Wen, Z.-L. (2012). Live Imaging Reveals Differing Roles of Macrophages and Neutrophils during Zebrafish Tail Fin Regeneration. Journal of biological chemistry. https://doi.org/10.1074/jbc.M112.349126

Li, V. S., S., SS, N., PJ, B., TY, L., WR, K., JP, G., . . . H, C. (2012). Wnt signaling through inhibition of β-catenin degradation in an intact Axin1 complex. Cell, 149(6). https://doi.org/10.1016/j.cell.2012.05.002

Liang, J., Wang, D., Renaud, G., Wolfsberg, T. G., Wilson, A. F., & Burgess, S. M. (2012). The stat3/socs3a pathway is a key regulator of hair cell regeneration in zebrafish. [corrected]. J Neurosci, 32(31), 10662–10673. https://doi.org/10.1523/jneurosci.5785-10.2012

Liu, K., Jiang, M., Lu, Y., Chen, H., Sun, J., Wu, S., . . . Que, J. (2013). Sox2 Cooperates with Inflammation-Mediated Stat3 Activation in the Malignant Transformation of Foregut Basal Progenitor Cells. Cell Stem Cell, 12(3), 304–315. https://doi.org/10.1016/j.stem.2013.01.007

Lorchner, H., Poling, J., Gajawada, P., Hou, Y., Polyakova, V., Kostin, S., . . . Braun, T. (2015).Myocardial healing requires Reg3beta-dependent accumulation of macrophages in the ischemic heart. Nat Med, 21(4), 353–362. https://doi.org/10.1038/nm.3816

Lotan, M., & Schwartz, M. (1994). Cross talk between the immune system and the nervous system in response to injury: implications for regeneration. FASEB J, 8(13), 1026–1033.

MacDonald, B. T., K, T., & X, H. (2009). Wnt/beta-catenin signaling: components, mechanisms, and diseases. Developmental Cell, 17(1). https://doi.org/10.1016/j.devcel.2009.06.016

Marjoram, L., Alvers, A., Deerhake, M. E., Bagwell, J., Mankiewicz, J., Cocchiaro, J. L., . . . Bagnat,M. (2015). Epigenetic control of intestinal barrier function and inflammation in zebrafish. PNAS. https://doi.org/10.1073/pnas.1424089112

Martinez, F. O., & Gordon, S. (2014). The M1 and M2 paradigm of macrophage activation: time for reassessment. F1000Prime Rep, *6*, 13. https://doi.org/10.12703/p6-13

Ming, G. L., & Song, H. (2011). Adult neurogenesis in the mammalian brain: significant answers and significant questions. Neuron, 70(4), 687–702. https://doi.org/10.1016/j.neuron.2011.05.001

Monje, M. L., Toda, H., & Palmer, T. D. (2003). Inflammatory blockade restores adult hippocampal neurogenesis. Science, 302(5651), 1760–1765. https://doi.org/10.1126/science.1088417

Mori, T., Miyamoto, T., Yoshida, H., Asakawa, M., Kawasumi, M., Kobayashi, T., . . . Yoshimura,A. (2011). IL-1beta and TNFalpha-initiated IL-6-STAT3 pathway is critical in mediatinginflammatory cytokines and RANKL expression in inflammatory arthritis. Int Immunol, 23(11), 701–712. https://doi.org/10.1093/intimm/dxr077

Nelson, C. M., Ackerman, K. M., O’Hayer, P., Bailey, T. J., Gorsuch, R. A., & Hyde, D. R. (2013). Tumor necrosis factor-alpha is produced by dying retinal neurons and is required for Muller glia proliferation during zebrafish retinal regeneration. J Neurosci, 33(15), 6524–6539. https://doi.org/10.1523/jneurosci.3838-12.2013

Nelson, C. M., Gorsuch, R. A., Bailey, T. J., Ackerman, K. M., Kassen, S. C., & Hyde, D. R. (2012).Stat3 defines three populations of Muller glia and is required for initiating maximal muller glia proliferation in the regenerating zebrafish retina. J Comp Neurol, 520(18), 4294–4311. https://doi.org/10.1002/cne.23213

Nguyen-Chi, M., Laplace-Builhe, B., Travnickova, J., Luz-Crawford, P., Tejedor, G., Lutfalla, G., . . .Djouad, F. (2017). TNF signaling and macrophages govern fin regeneration in zebrafish larvae. Cell Death Dis, 8(8), e2979. https://doi.org/10.1038/cddis.2017.374

Nguyen-Chi, M., Laplace-Builhé, B., Travnickova, J., Luz-Crawford, P., Tejedor, G., Lutfalla, G., . . .Djouad, F. (2017). TNF signaling and macrophages govern fin regeneration in zebrafish larvae [Research]. Cell Death & Disease, 8(8). https://doi.org/doi:10.1038/cddis.2017.374

Oosterhof, N., Kuil, L. E., van der Linde, H. C., Burm, S. M., Berdowski, W., van Ijcken, W. F. J., . .. van Ham, T. J. (2018). Colony-Stimulating Factor 1 Receptor (CSF1R) Regulates Microglia Density and Distribution, but Not Microglia Differentiation In Vivo. Cell Rep, 24(5), 1203–1217.e1206. https://doi.org/10.1016/j.celrep.2018.06.113

Paredes, M. F., Sorrells, S. F., Garcia-Verdugo, J. M., & Alvarez-Buylla, A. (2015). Brain size and limits to adult neurogenesis. J Comp Neurol. https://doi.org/10.1002/cne.23896

Parent, J. M. (2003). Injury-induced neurogenesis in the adult mammalian brain. Neuroscientist, 9(4), 261–272.

Parent, J. M., Valentin, V. V., & Lowenstein, D. H. (2002). Prolonged seizures increase proliferating neuroblasts in the adult rat subventricular zone-olfactory bulb pathway. J Neurosci, 22(8), 3174–3188. https://doi.org/20026296

Park, H. C., Shin, J., Roberts, R. K., & Appel, B. (2007). An olig2 reporter gene marks oligodendrocyte precursors in the postembryonic spinal cord of zebrafish. Dev Dyn, 236(12), 3402–3407. https://doi.org/10.1002/dvdy.21365

Peretto, P., & Bonfanti, L. (2014). Major unsolved points in adult neurogenesis: doors open on a translational future? Front Neurosci, 8. https://doi.org/10.3389/fnins.2014.00154

Petrie, T. A., Strand, N. S., Yang, C. T., Rabinowitz, J. S., & Moon, R. T. (2014). Macrophages modulate adult zebrafish tail fin regeneration. Development, 141(13), 2581–2591. https://doi.org/10.1242/dev.098459

Pilz, G. A., Bottes, S., Betizeau, M., Jorg, D. J., Carta, S., Simons, B. D., . . . Jessberger, S. (2018). Live imaging of neurogenesis in the adult mouse hippocampus. Science, 359(6376), 658–662. https://doi.org/10.1126/science.aao5056

Poss, K. D., Wilson, L. G., & Keating, M. T. (2002). Heart regeneration in zebrafish. Science, 298(5601), 2188–2190. https://doi.org/10.1126/science.1077857

Ramachandran, R., Reifler, A., Parent, J. M., & Goldman, D. (2010). Conditional gene expression and lineage tracing of tuba1a expressing cells during zebrafish development and retina regeneration. J Comp Neurol, 518(20), 4196–4212. https://doi.org/10.1002/cne.22448

Ramachandran, R., Zhao, X. F., & Goldman, D. (2011). Ascl1a/Dkk/beta-catenin signaling pathway is necessary and glycogen synthase kinase-3beta inhibition is sufficient for zebrafish retina regeneration. Proc Natl Acad Sci U S A, 108(38), 15858–15863. https://doi.org/10.1073/pnas.1107220108

Ransohoff, R. M. (2016). A polarizing question: do M1 and M2 microglia exist? Nat Neurosci, 19(8), 987–991. https://doi.org/10.1038/nn.4338

Ransohoff, R. M., & Brown, M. A. (2012). Innate immunity in the central nervous system. J Clin Invest, 122(4), 1164–1171. https://doi.org/10.1172/jci58644

Richmond, C. A., Rickner, H., Shah, M. S., Ediger, T., Deary, L., Zhou, F., . . . Breault, D. T. (2018). JAK/STAT-1 Signaling Is Required for Reserve Intestinal Stem Cell Activation during Intestinal Regeneration Following Acute Inflammation. Stem Cell Reports, 10(1), 17–26. https://doi.org/10.1016/j.stemcr.2017.11.015

Roehl, H. H. (2018). Linking wound response and inflammation to regeneration in the zebrafish larval fin. Int J Dev Biol, 62<otherinfo>(6-7-8), 473–477. https://doi.org/10.1387/ijdb.170331hr</otherinfo>

Sanz-Morejon, A., Garcia-Redondo, A. B., Reuter, H., Marques, I. J., Bates, T., Galardi-Castilla, M., . . . Mercader, N. (2019). Wilms Tumor 1b Expression Defines a Pro-regenerative Macrophage Subtype and Is Required for Organ Regeneration in the Zebrafish. Cell Rep, 28(5), 1296–1306.e1296. https://doi.org/10.1016/j.celrep.2019.06.091

Schindelin, J., Arganda-Carreras, I., Frise, E., Kaynig, V., Longair, M., Pietzsch, T., . . . Cardona, A. (2012). Fiji: an open-source platform for biological-image analysis. Nat Methods, 9(7), 676–682. https://doi.org/10.1038/nmeth.2019

Sehring, I. M., Jahn, C., & Weidinger, G. (2016). Zebrafish fin and heart: what’s special about regeneration? Curr Opin Genet Dev, 40, 48–56. https://doi.org/10.1016/j.gde.2016.05.011

Shiau, C. E., Kaufman, Z., Meireles, A. M., & Talbot, W. S. (2015). Differential requirement for irf8 in formation of embryonic and adult macrophages in zebrafish. PLoS One, 10(1), e0117513. https://doi.org/10.1371/journal.pone.0117513

Shigemoto-Mogami, Y., Hoshikawa, K., Goldman, J. E., Sekino, Y., & Sato, K. (2014). Microglia enhance neurogenesis and oligodendrogenesis in the early postnatal subventricular zone. J Neurosci, 34(6), 2231–2243. https://doi.org/10.1523/jneurosci.1619-13.2014

Shimizu, N., Kawakami, K., & Ishitani, T. (2012). Visualization and exploration of Tcf/Lef function using a highly responsive Wnt/beta-catenin signaling-reporter transgenic zebrafish. Dev Biol, 370(1), 71–85. https://doi.org/10.1016/j.ydbio.2012.07.016

Shimizu, Y., Ueda, Y., & Ohshima, T. (2018). Wnt signaling regulates proliferation and differentiation of radial glia in regenerative processes after stab injury in the optic tectum of adult zebrafish. Glia. https://doi.org/10.1002/glia.23311

Sierra, A., Encinas, J. M., Deudero, J. J., Chancey, J. H., Enikolopov, G., Overstreet-Wadiche, L. S., . . . Maletic-Savatic, M. (2010). Microglia shape adult hippocampal neurogenesis through apoptosis-coupled phagocytosis. Cell Stem Cell, 7(4), 483–495. https://doi.org/10.1016/j.stem.2010.08.014

Silver, J., & Miller, J. H. (2004). Regeneration beyond the glial scar. Nature Reviews Neuroscience, 5(2), 146–156. https://doi.org/doi:10.1038/nrn1326

Skaggs, K., Goldman, D., & Parent, J. M. (2014). Excitotoxic brain injury in adult zebrafish stimulates neurogenesis and long-distance neuronal integration. Glia, 62(12), 2061–2079. https://doi.org/10.1002/glia.22726

Sorrells, S. F., Paredes, M. F., Cebrian-Silla, A., Sandoval, K., Qi, D., Kelley, K. W., . . . Alvarez-Buylla, A. (2018). Human hippocampal neurogenesis drops sharply in children to undetectable levels in adults [Research]. Nature. https://doi.org/doi:10.1038/nature25975

Sun, F., Park, K. K., Belin, S., Wang, D., Lu, T., Chen, G., . . . He, Z. (2011). Sustained axon regeneration induced by co-deletion of PTEN and SOCS3. Nature, 480(7377), 372–375. https://doi.org/10.1038/nature10594

Than-Trong, E., & Bally-Cuif, L. (2015). Radial glia and neural progenitors in the adult zebrafish central nervous system. Glia, 63(8), 1406–1428. https://doi.org/10.1002/glia.22856

Todd, L., Squires, N., Suarez, L., & Fischer, A. J. (2016). Jak/Stat signaling regulates the proliferation and neurogenic potential of Muller glia-derived progenitor cells in the avian retina. Sci Rep, 6, 35703. https://doi.org/10.1038/srep35703

Tsarouchas, T. M., Wehner, D., Cavone, L., Munir, T., Keatinge, M., Lambertus, M., . . . Becker, C. G. (2018). Dynamic control of proinflammatory cytokines Il-1β and Tnf-α by macrophages is necessary for functional spinal cord regeneration in zebrafish. bioRxiv. https://doi.org/10.1101/332197

Tsarouchas, T. M., Wehner, D., Cavone, L., Munir, T., Keatinge, M., Lambertus, M., . . . Becker, C. G. (2018). Dynamic control of proinflammatory cytokines Il-1beta and Tnf-alpha by macrophages in zebrafish spinal cord regeneration. Nat Commun, 9(1), 4670. https://doi.org/10.1038/s41467-018-07036-w

Unger, M. S., Schernthaner, P., Marschallinger, J., Mrowetz, H., & Aigner, L. (2018). Microglia prevent peripheral immune cell invasion and promote an anti-inflammatory environment in the brain of APP-PS1 transgenic mice. J Neuroinflammation, 15(1), 274. https://doi.org/10.1186/s12974-018-1304-4

Walton, E. M., Cronan, M. R., Beerman, R. W., & Tobin, D. M. (2015). The Macrophage-Specific Promoter mfap4 Allows Live, Long-Term Analysis of Macrophage Behavior during Mycobacterial Infection in Zebrafish. PLoS One, 10(10), e0138949. https://doi.org/10.1371/journal.pone.0138949

Wan, J., Ramachandran, R., & Goldman, D. (2012). HB-EGF is necessary and sufficient for Muller glia dedifferentiation and retina regeneration. Dev Cell, 22(2), 334–347. https://doi.org/10.1016/j.devcel.2011.11.020

Wan, J., Zhao, X. F., Vojtek, A., & Goldman, D. (2014). Retinal injury, growth factors, and cytokines converge on β-catenin and pStat3 signaling to stimulate retina regeneration. Cell Rep, 9(1), 285–297. https://doi.org/10.1016/j.celrep.2014.08.048

Wang, X., Chen, H., Tian, R., Zhang, Y., Drutskaya, M. S., Wang, C., . . . Jiang, Y. (2017). Macrophages induce AKT/beta-catenin-dependent Lgr5(+) stem cell activation and hair follicle regeneration through TNF. Nat Commun, 8, 14091. https://doi.org/10.1038/ncomms14091

Wehner, D., Cizelsky, W., Vasudevaro, M. D., Ozhan, G., Haase, C., Kagermeier-Schenk, B., . . . Weidinger, G. (2014). Wnt/beta-catenin signaling defines organizing centers that orchestrate growth and differentiation of the regenerating zebrafish caudal fin. Cell Rep, 6(3), 467–481. https://doi.org/10.1016/j.celrep.2013.12.036

Zambusi, A., & Ninkovic, J. (2020). Regeneration of the central nervous system-principles from brain regeneration in adult zebrafish. World J Stem Cells, 12(1), 8–24. https://doi.org/10.4252/wjsc.v12.i1.8

Zhao, X. F., & Goldman, D. (2014). A new transgenic line reporting pStat3 signaling in glia. Zebrafish, 11(6), 588–589. https://doi.org/10.1089/zeb.2014.1502

Zhao, X. F., Wan, J., Powell, C., Ramachandran, R., Myers, M. G., & Goldman, D. (2014). Leptin and IL-6 family cytokines synergize to stimulate Müller glia reprogramming and retina regeneration. Cell Rep, 9(1), 272–284. https://doi.org/10.1016/j.celrep.2014.08.047

Zupanc, G. K., & Sirbulescu, R. F. (2011). Adult neurogenesis and neuronal regeneration in the central nervous system of teleost fish. Eur J Neurosci, 34(6), 917–929. https://doi.org/10.1111/j.1460-9568.2011.07854.x

